# Rapid, interpretable, data-driven models of neural dynamics using Recurrent Mechanistic Models

**DOI:** 10.1101/2024.10.10.617633

**Authors:** Thiago B. Burghi, Maria Ivanova, Ekaterina Morozova, Huaxinyu Wang, Eve Marder, Timothy O’Leary

**Affiliations:** Department of Engineering, University of Cambridge, United Kingdom; Department of Biology, Brandeis University, Waltham MA, USA 02543

**Author notes:** Author contributions: T.B. Burghi designed the model architecture and corresponding training algorithms, performed theoretical analyses, wrote the computer code implementation of the method, helped with electrophysiology experiments, and wrote the main body of the manuscript. M. Ivanova and E. Morozova performed the electrophysiology experiments, wrote the corresponding Methods subsection, and provided feedback on the manuscript. H. Wang helped with the computer code implementation of the training algorithms. E. Marder contributed to experiment design, experimental facilities, project supervision, and manuscript feedback. T. O’Leary conceptualized the project, contributed to experiment design, and helped writing the main body of the manuscript.

**Keywords:** Biophysical models, Deep learning, Electrophysiology, Central Pattern Generators, Neural Dynamics

## Abstract

Obtaining predictive models of a neural system is notoriously challenging. Detailed models suffer from excess model complexity and are difficult to fit efficiently. Simplified models must negotiate a tradeoff between tractability, predictive power and ease of interpretation. We present a new modelling paradigm for estimating predictive mechanistic-like models of neurons and small circuits that navigates these issues using methods from systems theory. The key insight is that membrane currents can be modelled using two scalable system components optimized for learning: linear state space models, and nonlinear artificial neural networks (ANNs). Combining these components, we construct two types of membrane currents: lumped currents, which are flexible, and data-driven conductance-based currents, which are interpretable. The resulting class of models — which we call Recurrent Mechanistic Models (RMMs) — can be trained in a matter of seconds to minutes on intracellular recordings during an electrophysiology experiment, representing a step change in performance over previous approaches. As a proof-of-principle, we use RMMs to learn the dynamics of two groups of neurons, and their synaptic connections, in the Stomatogastric Ganglion (STG), a well-known central pattern generator. We show that RMMs are efficiently trained using teacher forcing and multiple shooting. Due to their reliability, efficiency and interpretability, RMMs enable qualitatively new kinds of experiments using predictive models in closed-loop neurophysiology and online estimation of neural properties in living preparations.

**Significance Statement:** Our ability to understand the nervous system has been hindered by the difficulty of constructing good predictive models of neurons and circuits. This difficulty persists despite vast accumulated knowledge of how the basic components of the nervous system work, marking a gap in our ability to explain neural dynamics in terms of underlying mechanisms. This work describes a new data-driven modelling approach for neurons and small circuits that combines predictive power with mechanistic interpretation.

---

A major obstacle to progress in neuroscience is the difficulty of reconciling models with data. This difficulty is particularly acute in neural circuits that have non-trivial intrinsic dynamics, where the interplay of membrane currents over many timescales dictate circuit-level properties. Biophysically detailed models are fully interpretable, but suffer from excessive complexity and are notoriously difficult to fit to data (1–10). On the other hand, purely data-driven models may fit data very well, but their general purpose structure limits mechanistic insights (11–18), and they can give spurious predictions outside of the strict domain of the data they are trained with. At the heart of these challenges is a paradox: the basic biophysical principles of neurons and networks are very well understood, yet the models we derive from such principles struggle to predict neural activity as well as their phenomenological counterparts (19). On top of this paradox, most existing methods share the common drawback of being costly to constrain in terms of data, computational resources, and time.

In this work we present a new, efficient data-driven modelling approach that navigates the middle ground between mechanistic detail and black-box models from machine learning. Our ultimate goal is to reliably estimate mechanistic neuronal behaviour during the time frame of an experiment. This goal is both pragmatic and scientific: obtaining a predictive model during the lifetime of an experiment opens the door to real-time estimation and closed-loop control of a living neural circuit. Secondly, this goal provides a strong test of our modelling assumptions, and of the efficiency and robustness of model training.

It is important to recognise existing approaches that can successfully fit experimental data and predict neural dynamics in certain settings, and contrast these with our approach and requirements. Existing approaches comprise two broad classes. The first assumes relatively simple neural dynamics and uses phenomenological models to predict spiking, to remarkable effect in some cases (1, 11, 12, 15). However, this approach cannot account for rich intrinsic dynamics seen in circuits such as central pattern generators. Secondly, phenomenological models do not account for mechanistic effects of membrane conductances, limiting interpretability and preventing their use in closed loop, predictive control experiments with dynamic clamp and neuromodulators. The second class of approaches attempts to constrain large, detailed models to data. This requires extremely precise measurements, “clean” preparations, detailed knowledge about a given neuron’s ion channel types, and large computational resources. Importantly, such approaches are not yet practical to use during the lifetime of an experiment where neuronal dynamics are uncertain (2–8, 10, 20). Furthermore, it is not clear that these models capture ‘true’ parameters as opposed to plausible parameter subspaces of mis-specified models (21), which raises questions about their parsimony, interpretability and predictive power.

To address these difficulties, we introduce Recurrent Mechanistic Models (RMMs), a novel modelling framework that leverages the predictive power of machine learning while retaining biophysical, mechanistic intuition. RMMs, which can be framed as class of recurrent neural networks (RNNs) with biophysically motivated constraints, use a combination of linear time-invariant (LTI) state space models and artificial neural networks (ANNs) to capture the dynamics of intrinsic and synaptic ionic currents. Exploiting this model structure, we show that RMMs can be primed for different purposes in which different levels of interpretability are required. RMMs can predict the trajectories of ionic conductances when a detailed model is warranted, effectively acting as a quickly-trainable surrogate for conductance-based models. In more general cases, including most experimental settings where specific ionic currents are unknown or uncertain, we show that fully data-driven lumped currents can be used to predict the dynamics of interconnected neurons and the synaptic current between them; we also show that data-driven lumped and mechanistic currents can be combined to obtain interpretable models in a challenging experimental context.

A key advantage of RMMs is that they can be reliably trained in a matter of seconds to minutes in a consumer-grade desktop computer, enabling their use during an electrophysiology experiment. Rapid training is achieved with teacher forcing (22), which allows RMMs to be trained in a non-recurrent fashion that exploits massive parallelization. We show how to improve on teacher forcing results using a recurrent method, multiple shooting (for a technical exposition, see (23)), that can be tuned so as to trade off speed of training for prediction accuracy, while avoiding the well-known problem of exploding gradients (24).

To provide a realistic experimental test of this methodology, we estimate the dynamics of coupled neurons in the Stomatogastric Ganglion, a well-studied Central Pattern Generator (CPG) in the crab *Cancer borealis*. This circuit provides an ideal testbed for our application, as the dynamics of its neurons are complex and its function is dictated by the action of neuromodulators, which shape membrane conductance properties. We find that RMMs provide excellent predictive power during the lifetime of an experiment, and can recover features of underlying membrane conductances, enabling mechanistic interpretation.

## Results

Our results introduce Recurrent Mechanistic Models (RMMs), and demonstrate that they can reliably infer and interpret complex neuronal dynamics, while also being fast enough to generate predictions during electrophysiological experiments.

### Recurrent Mechanistic Models

RMMs are set apart from other types of neuron models due to the use of linear time-invariant (LTI) state space systems in conjunction with artificial neural networks (ANNs) to model the ionic currents flowing through the neuronal membrane. The basic idea is illustrated in Figure 1 A: a state space system produces a set of dynamical features of the membrane voltage signal, and the features are recombined by a readout function, constructed with ANNs, to yield the membrane current. The state-space systems can be systematically designed to have orthogonal convolution kernels, resulting in dynamically rich features for the ANNs to recombine (see Methods). As the results will show, modelling flexibility can be traded off for interpretability by imposing architectural constraints on the readout function. A schematic of a RMM for a neuron with one synaptic input is shown in Figure 1 B. It is straightforward to recognise that the schematic is equivalent to the circuit diagram of a single-compartment conductance-based model (25), with intrinsic, synaptic, leak and applied currents summed in parallel and integrated to compute membrane voltage.

**Fig. 1.**
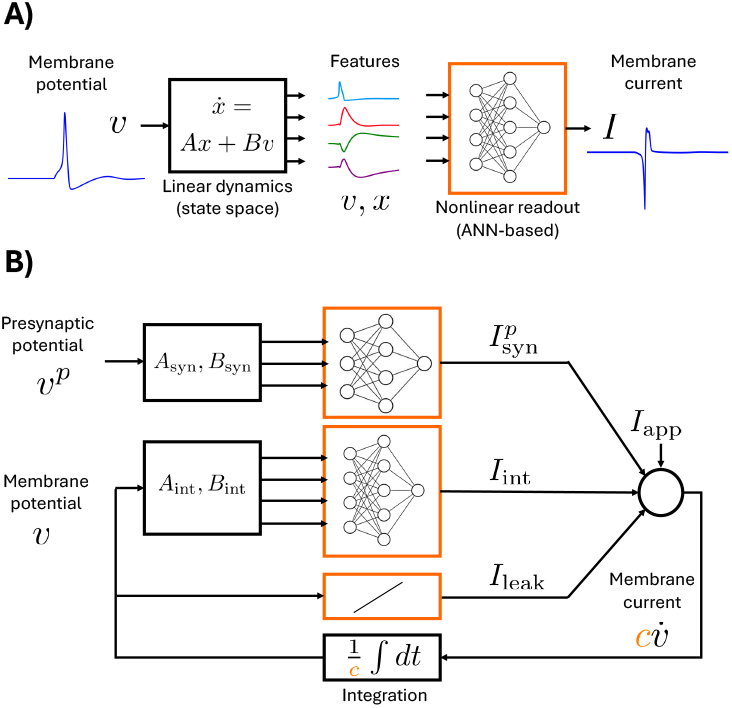
The basic structure of Recurrent Mechanistic Models. **A)** A state space system in series with an artificial neural network is used to transform membrane voltages into membrane currents. State-space systems are systematically designed so that features are dynamically rich (see Methods). Modelling flexibility can be traded off for interpretability by imposing architectural constraints on the ANN. **B)** A neuronal model is obtained by integrating applied, intrinsic and synaptic membrane currents. The blocks depicted in orange are those containing the main trainable parameters of the model, which also include the membrane capacitance, *c*. While state space models can in principle be learned, biophysical intuition can be used to construct them and keep them fixed, leading to faster training times (see Methods). Dots indicate time derivatives; in practice, the model is implemented in discrete-time according to Eq. (1)-Eq. (6).

We now briefly describe the basic mathematical structure of RMMs; further details can be found in Methods. Since RMMs are learned from sampled data, we use discrete-time notation. The discretized derivative voltage derivative is denoted by

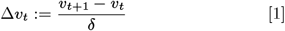

where *δ* is the sampling period used when collecting data in an experiment. The dynamics of a single neuron in our modelling paradigm is given by

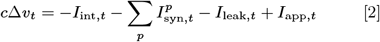

where, at time index *t*, the current *I*_int,*t*_ denotes the total intrinsic (ionic) current flowing through the neuron’s membrane, 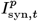 denotes the total synaptic current due to the *p*^th^ presynaptic neuron, *I*_leak,*t*_ denotes the leak current, and *I*_app,*t*_ denotes the applied current.

The total intrinsic current *I*_int,*t*_ models the sum of all ionic currents flowing through the neuron’s membrane. In general, the intrinsic current is given by

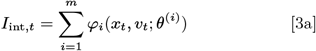

where each *φ*_*i*_ is an ANN-based nonlinear readout function, parameterized by *θ*^(*i*)^, and *x*_*t*_ is the vector-valued state of a linear time-invariant system given by

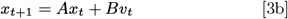

with *A* and *B* matrices of the appropriate dimension. Although *A* and *B* can in principle be learned, we have empirically found a systematic choice for *A* and *B* such that state space parameters do not need to be learned to achieve high predictive performance. This is described in *constructing linear systems* in Methods.

Each learnable ANN-based readout *φ*_*i*_ in Eq. (3a) models an individual component of the total intrinsic current, whose level of mechanistic detail is determined by the parametric form of *φ*_*i*_. To model electrical current components whose mechanisms are unknown, uncertain, or too complex to deduce, we employ **lumped currents**. Their models do not single out individual ionic types, and are constructed with a multi-layer perceptron (MLP, i.e, a black-box feed-forward ANN) according to

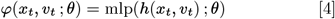

where mlp(·; *θ*) is a single-output MLP, and *h*(*x*_*t*_, *v*_*t*_) is a fixed layer used to normalize and possibly reduce the dimension of the state space output vector 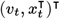. To model electrical current components whose ionic mechanisms are better understood, we employ **data-driven mechanistic currents**. Their models make use of mechanistic priors, such as the reversal potential, or the type of gating dynamics of a current (activation, inactivation, or both). A typical data-driven conductance-based current is given by

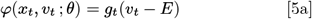

with

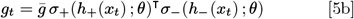

where *E* is the reversal potential and 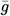 is a conductance scaling parameter (about which we might have prior knowledge). Here, *σ*_+_(·; *θ*) and *σ*_−_(·; *θ*) are single-layer perceptrons whose weight matrices are constrained so as to model activation (+) and inactivation (−) gating variables, respectively. For more details on the models given by Eq. (3b)-Eq. (5), see *constructing lumped currents* and *constructing mechanistic currents* in Methods.

The justification for this particular model structure is based on practical as well as theoretical considerations. From a pragmatic viewpoint, using a single state space system to model the recurrent part of the intrinsic current dynamics greatly facilitates the choice of initial model parameters, which reduce to a sequence of time constants (Methods). State space models also have a smaller number of parameters when compared to equivalent LTI representations such as temporal convolutions. Finally, the structure of RMMs allows exploiting massive paralellization during training, since the readouts *φ*_*i*_ of every intrinsic current component share the same “reservoir” of internal states *x*_*t*_ (see *training algorithms* in Methods).

From the theoretical point of view, we are supported by the fact that any reasonable model of an ionic current (or conductance) must have the so-called Fading Memory property (27). This means that the mapping from voltage to current (or conductance) can be approximated arbitrarily well by a so-called Wiener model: a LTI system with a nonlinear static readout function (27, 28). As pointed out in (13), fading memory can be empirically observed in the input-output behaviour of ionic currents under voltage-clamp. Intuitively, fading memory holds in a population of ionic currents if clamping the membrane voltage to a particular value results in a unique steady-state current, regardless of the value of the voltage prior to clamping. It is the very reason for which neuronal excitability can be studied using steady-state IV curves.

The total synaptic current 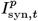 caused by a presynaptic neuron with membrane potential 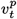 is modelled analogously to the intrinsic current, the only difference being the use of 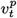 instead of *v*_*t*_ in Eq. (3b). Finally, the leak current is given by a linear-in-the-parameters model

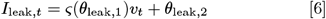

where the softplus function, defined by *ς*(·) = log(1 + exp(·)), is used to keep the leak conductance positive throughout training. The leak model Eq. (6) can also be re-parametrized in terms of maximal conductance and reversal potential.

### RMMs recover conductance trajectories from current-clamp data

The first question we ask is whether RMMs are able to predict the membrane voltage and conductance trajectories of conductance-based models, based solely on applied current– voltage data. To test this, we employed data-driven conductances (Eq. (5)) to design an RMM whose currents mimic the sodium and the potassium currents of the Hodgkin-Huxley neuron (26). The structure of the resulting total intrinsic current is illustrated in Figure 2 A, and the parameters are described in detail in *Hodgkin-Huxley RMM* in Methods. In constructing this model, we exploit *timescale separation*: the state space system is defined with four “fast” states evolving roughly in the timescales of sodium activation (0.1, 0.3, 0.5, and 0.7 milliseconds), and four “slow” states corresponding to sodium inactivation and potassium activation (1, 3, 5, and 7 milliseconds). While the fast state components are fed to the sodium activation readout, the slow state components are fed to both the sodium inactivation and the potassium inactivation readouts. In addition, the fixed layers at the input of the activation and inactivation readouts are designed so that, at randomly initialized weights and biases, both mechanistic currents are roughly inactivated outside of the range of voltages relevant to the HH model’s dynamics (see Methods).

**Fig. 2.**
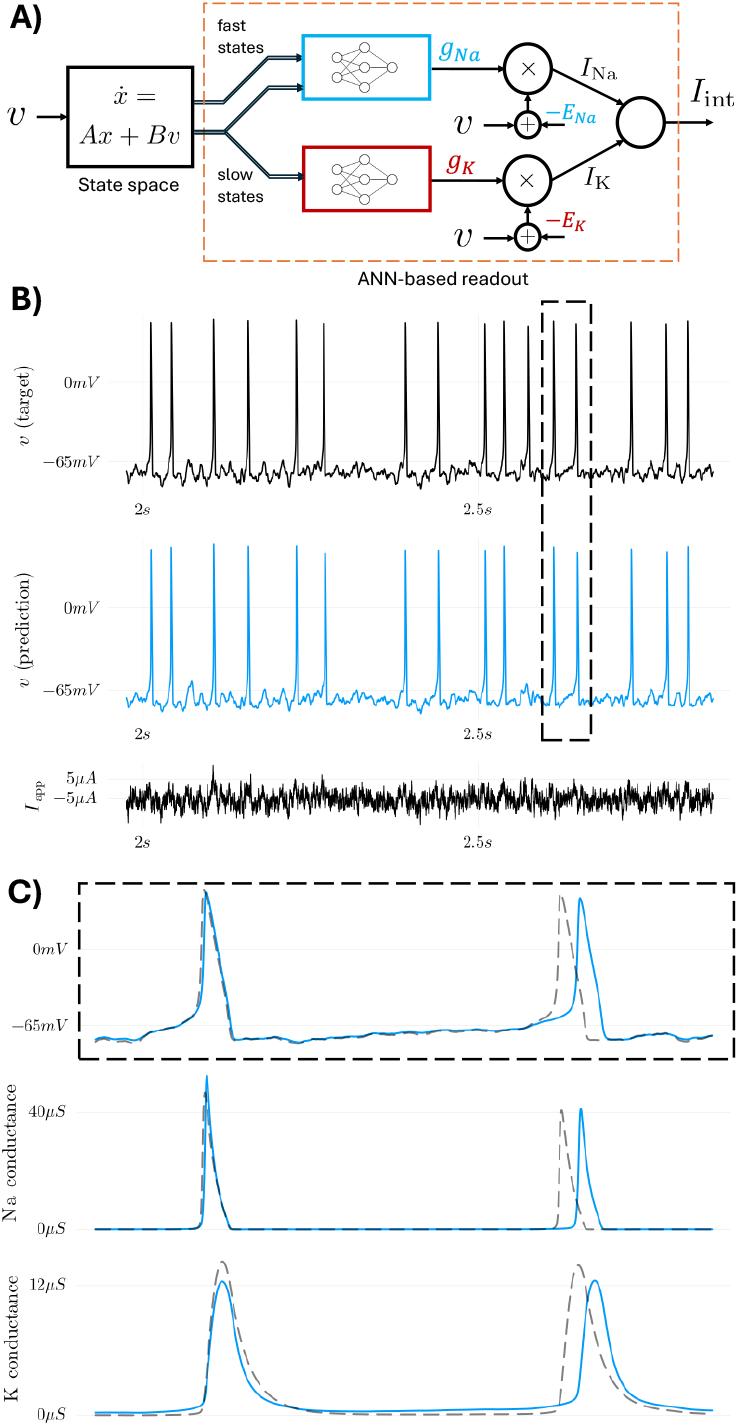
RMMs recover the classical Hodgkin-Huxley conductances (26) from input-output data. **A)** A state space system with eight internal states is designed so that the states evolve in fast and slow timescales roughly corresponding to those of the spiking dynamics of the original HH model. The ANN-based readout is constrained to promote interpretability (cf. Figure 1): sodium and potassium conductances are obtained as outputs of MLPs, and the corresponding ionic currents are obtained by multiplying those data-driven conductances by a difference in potential (*v − E*_ion_), with *E*_ion_ a prior reversal potential. To ensure the model learns the correct currents, fixed layers selecting fast and slow states are used. **B)** Target voltage trace (top) resulting from simulating the original HH model given a noisy applied current, and corresponding RMM prediction (bottom) on held-out test data. **C)** (top) zoom of the dashed region in (B) shows accuracy of spike prediction. (middle, bottom) The RMM accurately recovers conductance trajectories during spiking. For modelling and training details, see *Hodgkin-Huxley RMM* in Methods.

After training a randomly initialized Hodgkin-Huxley RMM with regularized teacher forcing (TF) on a current-clamp dataset, we simulated the trained RMM on held-out validation data, obtaining the the predictions shown in Figure 2 B and C (for a description of TF and the training dataset, see Methods). It can be seen that the learned RMM predicts not only the spike times of the target HH model, but also its ionic conductance trajectories. In Figure S1, we reproduce these results for five different RMMs initialized and trained with different random number generator (RNG) seeds chosen arbitrarily; all models are obtained around the same training epoch, and display similar predictive power at the level of spike times and conductance trajectory shapes. Notice that the small difference in the shape of conductance traces is an expected consequence of model mismatch, and of the fact that conductance traces are not used in the cost function of the training problem (which only uses applied current and voltage signals).

### Experimental validation of the RMM approach

While data-driven conductances (Eq. (5)) can make RMMs fully interpretable, in most experimental scenarios the precise ion channel makeup of a given neuron is not readily available, providing poor biophysical priors to work with. In addition, quantitatively predicting the response of neuronal circuits *in vitro* and *in vivo* often requires using multi-compartment models whose parameters are difficult if not impossible to estimate based on a single voltage signal. This section shows how single-compartment RMMs can address these problems through judicious use of lumped currents (Eq. (4)).

As a proof-of-principle that RMMs can be used to estimate the behaviour of neurons with complex morphology and uncertain biophysics *solely from applied current and voltage measurements*, we performed experiments to predict neuronal dynamics in a well-studied system: the Stomatogastric Ganglion (STG) of the crab *Cancer borealis*. We use the experimental setup illustrated in Figure 3 A: after pharmacologically blocking glutamatergic synapses of the STG, a simple feed-forward network is obtained where the Lateral Pyloric (LP) neuron is synaptically driven by two Pyloric Dilator (PD) neurons via cholinergic synapses. Both PD neurons are strongly coupled to the Anterior Burster (AB) neuron, a bursting pacemaker. As an approximation, one can think of this as a feedforward circuit where a strongly coupled subcircuit consisting of the two PD neurons and the AB neuron drives the single LP neuron. The only data used for training our models in this section consists of the applied currents and voltage signals from the LP neuron and one of the PD neurons. The ‘true’ PD-AB dynamics in particular are thus extremely complex, owing to the morphology of the cells as well as the fact that the PD measurement signal accounts for the combined dynamics of three coupled neurons.

**Fig. 3.**
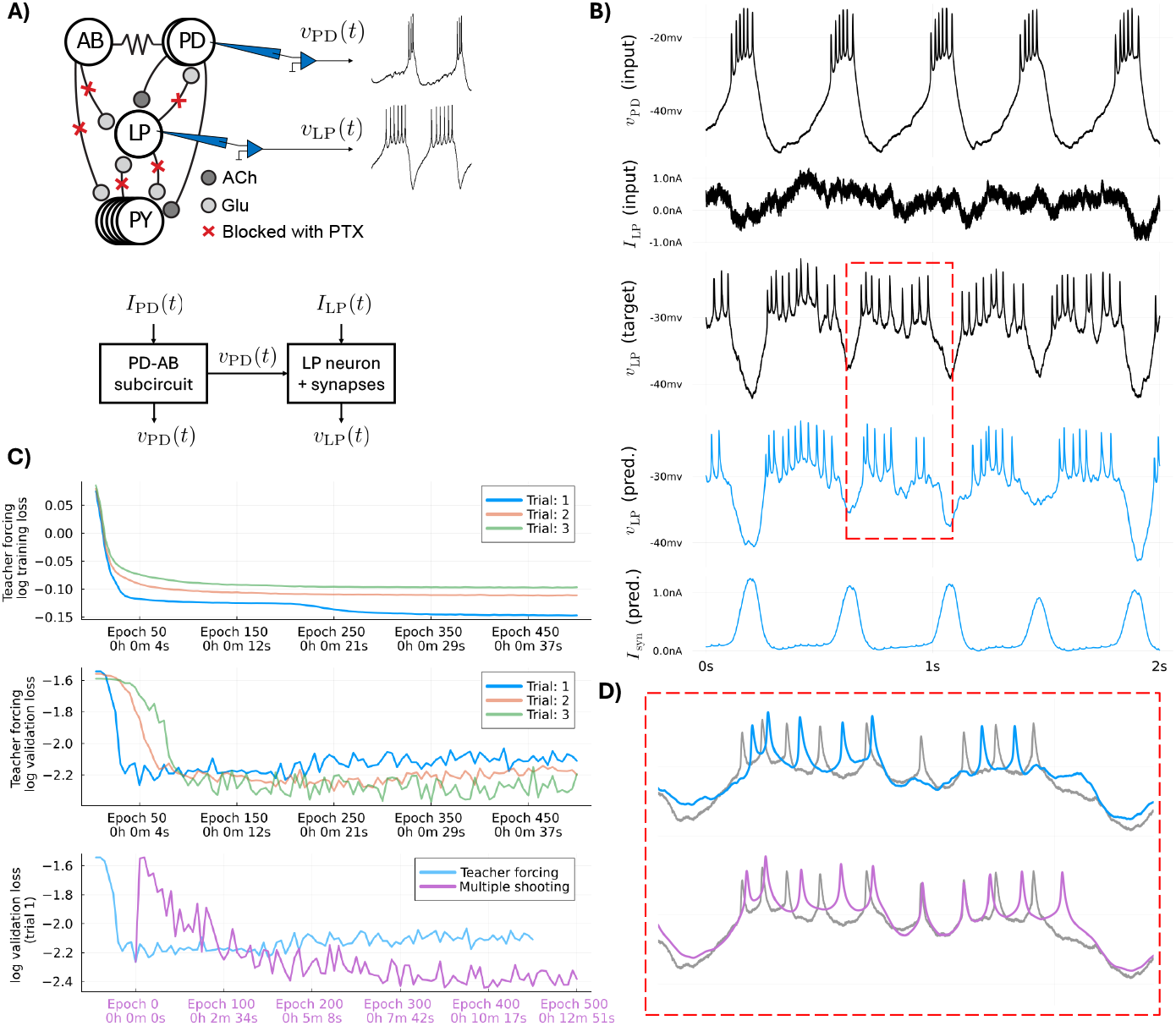
Estimating a RMM model of a neuron with complex spatiotemporal dynamics. **A)** Experimental setup used to generate data for fitting an RMM: the STG circuit is bathed in PTX, which blocks Glutamate synapses but not Acetylcholine synapses; intracellular recordings are taken of *one* of the PD neurons and the LP neuron; the resulting system is a feedforward interconnection with the measured presynaptic PD neuron voltage driving the LP neuron. **B)** Example voltage traces showing held-out data (three top panels) and the predicted voltage and synaptic currents from an RMM trained with teacher forcing (see Methods). **C)** Evolution of the loss functions during training for three different trial datasets; each dataset consists of around 2 seconds of data recorded with a sampling period of 0.1 ms. The highlighted (blue) Trial 1 yields the model used an example in (B); predictions of the models trained on Trials 2 and 3 are shown in Figure S2. Top: loss function value during training with teacher forcing, evaluated with held-out data; middle: validation loss function value using models trained with teacher forcing; bottom: multiple shooting improves the performance of the best model obtained with teacher forcing. **D)** Example traces demonstrating the improvement obtained with multiple shooting. For a description of teacher forcing and multiple shooting, as well as the loss function used for model validation, see Methods.

### Rapid prediction of total intrinsic and synaptic currents

To test the predictive power of RMMs constructed exclusively with lumped currents, we trained models to predict the responses of the LP neuron in Figure 3 A. The LP neuron RMM is constructed according to the diagram in Figure 1 B, with lumped ANN readouts given by Eq. (4) for both intrinsic and synaptic currents (for details, see *LP neuron RMM* in Methods). To generate a dynamically rich, time-varying input to LP, we injected noisy currents given by an Ornstein-Uhlenbeck process, while monitoring both LP and PD membrane potentials. After fitting an RMM we analysed the predictive performance of the model on held-out test data (Figure 3 B) and found good quantitative agreement between predicted LP membrane voltage (blue traces) and measured LP membrane voltage. The model also enabled us to infer the total synaptic current *I*_syn_, which varied as expected with PD bursts to generate intermittent pauses in LP spiking.

Over multiple fitting runs with validation on held-out data, we found that good steady-state prediction error is achieved independently of initial model parameters and datasets (Figure 3 C). In Figure 3 C (top), we plotted training losses for three different trials, corresponding to datasets collected from the same preparation at different times. Blue curves correspond to the model of Figure 3 B, while orange and green curves correspond to models whose predictive performance is showcased in Figure S2. Teacher forcing losses decrease and stabilize rapidly, under a minute. These results thus demonstrate a crucial feature of RMMs: the fact that reasonably good prediction accuracy, under large system uncertainty, can be achieved efficiently, with prediction error asymptoting in several tens of seconds on modest hardware (see Methods).

The learning performance of RMMs is a result of using an efficient training method based on teacher forcing (Methods). As with all machine learning methods, the decrease of training loss during teacher forcing does not guarantee a decrease in predictive power, as shown in Figure 3 (C, middle), where validation losses are plotted for “model snapshots” taken at different epochs during training (see Methods for a description of training vs. validation losses). Thus, while teacher forcing quickly trains the model, it also leads to overfitting, making early stopping or strong regularisation essential.

If necessary, predictive accuracy can be enhanced further using a generalisation of teacher forcing called multiple-shooting (see Methods). The result of this additional training on predictive accuracy is shown in Figure 3 (C, bottom), while Figure 3 D shows how spiking traces from the zoomed in portion of A are improved with this extra step of training. Starting from a model trained with teacher forcing, multiple shooting improves model performance at the cost of longer training times, up to several minutes on this dataset. The tradeoff between predictive accuracy and training time can therefore be dictated by the requirements of the experiment, and can yield very good results in minutes.

### Predicting and interpreting endogenous bursting

We next used the RMM framework to learn an interpretable single-compartment model of the bursting PD-AB subcircuit, often called the pacemaker kernel, in the same experimental setup of Figure 3 A. Mechanistically, bursting emerges from a complex combination of sodium, potassium, and calcium ion channels which are unevenly distributed in the system. Despite the complexity, modelling the pacemaker kernel with a single burst-excitable compartment is made possible by the strong electrical coupling between AB and PD neurons, which allows us to assume that the slow component of the bursting wave is roughly the same across those neuron’s membranes. However, quickly fitting mechanistic models of the pacemaker kernel remains difficult. In particular, the sodium channels driving intra-burst spikes are located in a region of the PD neuron’s axon far from the recording site in the soma, as illustrated in Figure 4 A. As a result, fast intra-burst spikes in the recordings are heavily filtered versions of the PD neuron’s spikes (see (29, 30) for details).

**Fig. 4.**
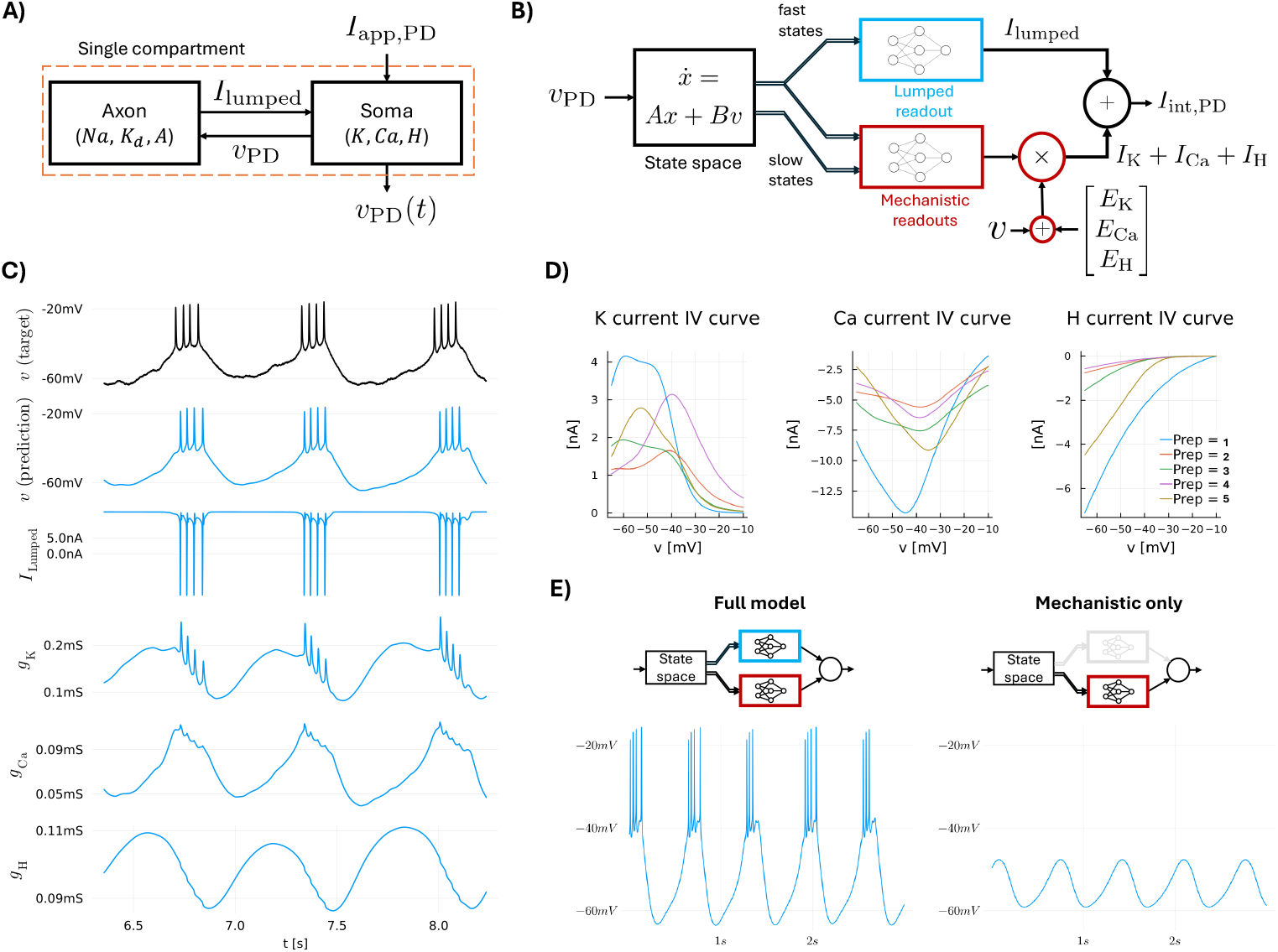
Training a RMM with lumped and mechanistic branches to learn endogeneous bursting rhythms. **A)** a diagram representing the dynamics of the AB-PD pacemaker, simplified here as a single PD cell (30). Fitting traditional mechanistic models to the system is challenging: a single-compartment conductance-based model cannot quantitatively predict the membrane voltage at the soma (where recordings are made), since fast sodium and potassium currents are located in a distant part of the axon. **B)** To quickly obtain a predictive single-compartment model, we employ an RMM with a mix of lumped and mechanistic currents. The lumped current, which does not distinguish between ion channels and does not require mechanistic structure, is used to predict the fast axonal current. To model the calcium and potassium channels of the soma, we employ three data-driven mechanistic currents to capture the aggregate dynamics of potassium, calcium, and hyperpolarization-induced currents. **C)** Validation of an RMM model on held-out membrane voltage data (black traces). The model, which was trained on current-clamp data (see *PD neuron RMM* in Methods) predicts the membrane voltage well. In addition, the model allows interpreting its internal variables as individual currents and conductances. Figure S5 shows similar results for other PD neuron preparations. **D)** Steady-state IV curves of four RMMs fit to different PD neuron preparations show that RMMs are able to learn the basic mechanisms of bursting based on calcium and potassium currents. **E)** Removing the fast lumped current from the RMM uncovers the slow wave dynamics underlying burst excitability. All models in this figure and Figure S5 were trained with teacher forcing (see methods) over the course of 200 epochs, corresponding to about two minutes of training (see Figure S6 for training curves).

To deal with this challenging scenario, we used a combination of lumped and mechanistic data-driven currents to learn the dynamics of the AB-PD pacemaker, as depicted in Figure 4 B, and asked whether the resulting RMM would yield the appropriate intrinsic mechanism for generating endogenous bursting. The RMM is constructed with three mechanistic currents representing the aggregate dynamics of calcium currents (*I*_Ca_), potassium (including calcium-activated) currents (*I*_K_), and hyperpolarization-induced currents (*I*_H_). To model the intra-burst spiking dynamics while constraining the RMM to a single compartment, a lumped current (*I*_lumped_) using fast state components is used; this current is more heavily regularized during training than mechanistic currents (see *PD neuron RMM* in Methods).

Figure 4 C, top, shows example voltage traces of a PD neuron recording. The voltage trace shown was held out from training and used to validate a pacemaker RMM, whose predictions are shown in blue in Figure 4 C. The RMM, which was trained in around two minutes (Figure S6), predicts the target dataset well. In addition to predicting the target voltage, the structure of the RMM allows recovering the lumped membrane current, representing the fast axonal current, as well as the data-driven conductances associated to the aggregate calcium, potassium, and H currents. With minimal a priori information, the RMM is able to learn the functional role of the three types of currents: the aggregate potassium current activates following a burst, hyperpolarizing the membrane; the aggregate calcium current drives the upstroke of the slow bursting wave; and the aggregate *H* current activates following hyperpolarization, helping to initiate the burst.

To evaluate the consistency of the RMM method, we fitted RMMs to four other PD neuron preparations, using the same hyperparameters as those used to obtain Figure 4 — except for the overall regularization constant *ρ*, which was tuned to deal with differing noise levels. The results are shown in Figure S5, where it can be seen that even though the recordings are done in the same type of cell, the features of the recorded voltage traces are reasonably different, due to the intrinsic variability between individual preparations. In Figure 4 D we plot the steady-state IV curves of the inferred aggregate potassium and calcium currents across all five preparations. The steady-state IV curves all overlap in the range usually associated to the activation of calcium currents in PD neurons of the STG, indicating self-consistency among the preparations, as well as consistency with known voltage-clamp properties of these currents form previous work. We note, that since the models were trained on current-clamp data alone, we would expect that the steady-state IV curves shown in Figure 4 D are subject to greater uncertainty than current-clamp voltage predictions, such as those in Figure 4 C (blue traces). However, these results indicate that RMMs can extract consistent, mechanistically relevant features of membrane physiology without the need for voltage-clamping.

To further investigate whether RMM predictions at the level of individual conductances and currents are dynamically meaningful and generalizable, we probed the behaviour of the RMM after removing the lumped axonal fast current from the model. Figure 4 E shows the result for the same model as Figure 4 C. In the left panel, it can be seen that with both lumped and mechanistic currents, and no external excitation, the RMM generates an endogenous bursting rhythm. This demonstrates generalization with respect to training data, as the model was not trained with a constant current. Removing the fast lumped current results in a slow wave, shown in the right panel, proving the point that the mechanistic currents alone are sufficient to drive the endogenous rhythm.

## Discussion

Unlike some areas of physical sciences, where quantitative predictions can determine the validity of a theory, model predictions in neuroscience often have a more equivocal character. A poor fit to data can imply incorrect modelling assumptions, low quality data, inappropriate fitting methods, or any of these issues simultaneously. This ambiguity persists even when modelling reduced preparations consisting of a small number of neurons, and where tight control of experimental conditions and accurate measurements are possible. These challenges have preoccupied neurophysiologists for decades, and the limited success in handling them has hindered the field’s ability to apply vast accumulated knowledge of low-level physiology to systems-level questions.

The approach we have introduced here does not solve all these problems, but it does permit theoretically grounded predictive models to be inferred during the course of an experiment, which drastically enhances our ability to solidify theories of neural circuit function, and to use these theories to practical ends. Our goal was to find a model that captures what we know about biophysics without assuming more than can be measured, and to find a means of constraining the model to produce good predictive accuracy on a timescale of minutes.

Achieving this goal forced us to sacrifice a modelling formalism that is often placed among the most successful theoretical developments in all of neuroscience: the detailed Hodgkin-Huxley form of conductance based modelling (26). This may seem tantamount to abandoning any semblance of ‘mechanism’ in our modelling approach. Most mechanistic models are conductance-based and strive to account for ion channel gating dynamics and even the spatial distribution of channels in complex morphologies. Such models are fundamental for understanding how low level biophysics shapes the dynamic properties of neurons (30–33) and in some cases constitute predictive models that fit data very well (10). However, in general, the empirical success of this modelling paradigm has been modest, especially when applied to more typical neural circuits that possess many tens of ionic conductance types, complex morphologies and physiology that hinders precise experimental measurements. Constraining the parameters of detailed models under such conditions is notoriously difficult (34).

In addition to the practical difficulty of parameter estimation in detailed models, there is a deeper, scientific problem with attempts to capture detail. Regardless of their detail, models likely omit unknown structural and biophysical properties that are impractical to measure (35). In other words, the strict form of a detailed conductance based model that supposedly describes a given type of neuron is almost certainly wrong, and the parts of it that are right might constrain the model’s dynamics in unwanted ways, and even introduce artefacts (36). This simultaneously impacts predictive power and interpretability: inferred parameter values of a misspecified model, however precise, may have no sensible physical meaning.

Our approach acknowledges our ignorance of the extant pool of conductances and synaptic inputs in a preparation, their spatial distribution, phosphorylation state, splice variants and the remaining plethora structural variation that is known to exist but cannot be measured. Instead, we only model what we can measure, representing this information in terms of known biophysics, such as early and late IV relationships, and, where possible, inward and outward current components. Ultimately these are the features of the underlying biophysics that dictate a neuron’s dynamics, and in this sense, they are the most parsimonious, mechanistic inference we can be confident of in a typical experimental setting.

Even though the experimental system we chose is well characterised, with a relatively simple connectivity, unobserved inputs and dynamics in the STG abound in the data. Although we blocked fast inhibitory transmission with PTX, there are numerous modulatory inputs coming from the front-end ganglia that are still present in the preparation that affect several intrinsic currents (37) in a manner that depends on the wider state of the preparation (38). Gap junctions between the membranes of different cells introduce further uncertainty (39). PD is tightly electrically coupled to AB, but it is also loosely coupled to LPG, which in turn is connected to VD, and several other cells. LP is connected to IC and multiple PY cells. Furthermore, the strength of gap junctions might be susceptible to modulation (40). These various inputs and perturbations are not accounted for in the model because we are unable to measure them directly, it is therefore far from trivial to be able to predict dynamics in this circuit with any degree of accuracy.

As with most models of the neuronal membrane, RMMs as presented in this paper assume that the applied current is injected at the site of membrane recordings. This assumption is informed by our choice of Discontinuous Current Clamp (DCC) as the method for current injection. DCC is practical, as it allows voltage to be measured simultaneously with the injection of applied current, using a single electrode. It is well-known that DCC results in significantly higher levels of measurement noise in the data. While improved results can in principle be obtained with two-electrode recordings and continuous-current clamp, in that case the effect of filtering of the fast components of the applied current by the neuronal membrane should be taken into account by the model.

## Methods

Results in this paper were obtained with code written in Julia and rely on the Flux.jl package (41). The code used to produce the results above can be found in https://github.com/thiagoburghi/RMM. In what follows we denote the all-ones and all-zeros vectors by **1** and **0**, respectively.

### Constructing linear systems

The state space matrices *A* and *B* in the total intrinsic current dynamics Eq. (3) are constructed systematically. First, we choose a sequence of *n* positive time constants *τ*_1_, *τ*_2_, …, *τ*_*n*_ believed to play a role in the collective dynamics of the intrinsic currents; second, we use the sequence {*τ*_*j*_} to construct a discrete-time state space system. The sequence {*τ*_*j*_} is most easily constructed by sampling the time constant functions of an existing “prior” conductance-based model^1^. Letting *τ*_*w*_ (*v*) denote the time constant function of a gating variable *w* in such a prior model, we set the smallest and largest time constants as

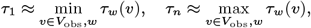

where *V*_obs_ = [*v*_min_, *v*_max_] is the observed membrane voltage range of the data. Intermediate time constants are chosen incrementally between *τ*_1_ and *τ*_*n*_ so as to roughly represent the range of values observed in all the *τ*_*w*_ (*v*). As an illustration of such time constant functions, in Figure S3 we plot the *τ*_*w*_ (*v*) of the LP neuron model of (42), which were used to inform the choices of {*τ*_*j*_} in this paper.

Once {*τ*_*j*_} is given, we compute a sequence {*λ*_*j*_} of eigenvalues for *A* (i.e., discrete-time system poles) by applying the zero-order-hold transformation

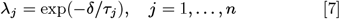

where *δ* is the sampling period of the data. This transformation leads to a stable state space system; see (43) for sampled-data system theory. The eigenvalues in Eq. (7) are then used to construct a *state space realization*: a choice of *A* and *B* such that the eigenvalues of *A* are given by *λ*_*j*_. This paper considers two types of realizations:

1. A *diagonal* realization where *A* = diag{*λ*_*j*_} is a diagonal matrix, and *B* = (**1** *−* vec{*λ*_*j*_)}.
2. A realization with *orthogonal impulse responses*^2^ (44). The SI Appendix describes how such a realization can be constructed from {*λ*_*j*_}.

An important fact about the two realizations above is that their DC (i.e., steady-state) gains (*I* − *A*)^*−*1^*B* are elementwise positive. Different realizations of *A* and *B* affect the learning algorithms by changing the trajectories of the inputs to ann-based readouts. For illustrative purposes, in Figure S4 we compare the “spike responses” of a diagonal state space system with those of a system with identical *A* matrix eigenvalues, but orthogonal impulse responses; the figure shows how the states of the latter overlap less in time, an effect we have found to be beneficial when training lumped currents. Intuitively, a system with orthogonal impulse responses generates a rich set of signals for ANNs to recombine.

### Constructing lumped currents

The MLPs used to model lumped readouts in Eq. (4) are given by the chained composition of mappings

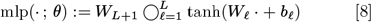

where *L* is the number of MLP layers, *W*_*𝓁*_ are weight matrices, *b*_*𝓁*_ are bias vectors, and tanh(·) applies the hyperbolic tangent elementwise to its inputs. In this work, the weight matrices are initialized with the Nguyen-Widrow heuristic (45), which is optimized for MLP inputs that lie in the interval [−1, 1] (see SI Appendix). The fixed layer *h*(*x*_*t*_, *v*_*t*_) of a lumped current is given by a non-learnable affine mapping used to normalize *v*_*t*_, as well as to select and normalize a subset of states in *x*_*t*_ which is passed as input to the MLP. In our results we have used min-max normalization, which maps the minimum and maximum values of the inputs of the MLP seen during training to −1 and +1, respectively (see Appendix SI for details). Lumped currents have empirically been found to work better with state space models with orthogonal impulse responses, see *LP neuron RMM* and *PD neuron RMM* below.

#### Constructing mechanistic currents

The *σ*_+_(·; *θ*) and *σ*_−_ (·; *θ*) mappings of mechanistic readouts in Eq. (5) are given by

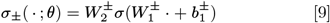

where *σ*(·) applies the logistic function (1 + exp(−·))^*−*1^ elementwise to its inputs. The sign of the elements of the weight matrices in Eq. (9) are constrained so as to implement monotonically increasing (*σ*_+_) and decreasing (*σ*_*−*_) mappings, in the sense that if *u*_2_ ≥ *u*_1_ elementwise, then *σ*_+_(*u*_2_) ≥ *σ*_+_(*u*_1_) and *σ* _−_ (*u*_2_) ≤ *σ* _−_ (*u*_1_), also elementwise. This is done to model the activation and the inactivation of a current, respectively. The weight constraints are implemented by setting 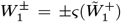 and 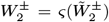, where *ς*(·) applies the softplus function log(1 +exp(·)) elementwise to its inputs. The matrices 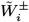 are unconstrained, and contain the trainable weights of a mechanistic readout; they are initialized randomly, but in such a way that

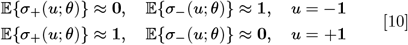

See the SI appendix for the detailed initialization heuristic.

The fixed layer *h*(*x*_*t*_) is a non-learnable affine normalisation mapping. It is used to select a subset of states of *x*_*t*_, and normalize those states. Normalisation implements a biophysical prior: based on a prior activation or inactivation voltage interval [*v*_low_, *v*_high_], *h*(*x*_*t*_) maps the steady-state values of *x*_*i,t*_ at *v*_low_ and at *v*_high_ to −1 and +1, respectively (SI Appendix). Notice that as a consequence of Eq. (10), this ensures the network is initialized in such a way that currents respect the prior activation/inactivation range [*v*_low_, *v*_high_].

Mechanistic readouts can read from either diagonal state space systems or systems with orthogonal impulse responses. But because of the constraints placed on the weight matrices of *σ*_±_ (·; *θ*), the choice of state space realisation is critical to the those currents. Reading from diagonal realisations results in “simple” currents where a monotonically increasing voltage trajectory leads to monotonically increasing activations *σ*_+_ and monotonically decreasing inactivations *σ*_*−*_. This is a straightforward fact from *positive systems theory* (46) (see details in the Appendix SI). Reading from realisations with orthogonal impulse responses results in more flexible (but also more degenerate) models, since in this case the preservation of the monotonicity of voltage trajectories is not guaranteed. Such realisations can be used in conjunction with mechanistic readouts to model complex ionic currents, such as calcium-activated potassium currents. Notice however the DC gain (*I*− *A*)^*−*1^*B* of such realisations is still elementwise positive. Hence, monotonically increasing/decreasing *steady-state voltages* lead to monotonically increasing/decreasing steady-state activations *σ*_+_, and monotonically decreasing/increasing steady-state inactivations *σ*_*−*_. This results in a biophysical constraint in the IV curves of the corresponding ionic currents.

In mechanistic currents, it is possible to gain further modelling flexibility by modifying Eq. (5) as

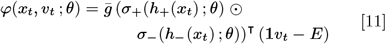

where ⊙ denotes the elementwise (Hadamard) product, and now *E* is a vector of Nernst potentials. This can be used when the Nernst potential of a current is uncertain, as is often the case with Calcium currents; the Nernst vector *E* acts as extra parameters of the data-driven mechanistic current.

#### Training RMMs

Rapid training is achieved with two related training methods that avoid well-known difficulties commonly encountered when training mechanistic models (47). We consider a dataset given by 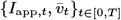 consisting of measured applied current *I*_app,*t*_ and voltage targets 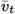. The main difficulty is that of so-called *exploding gradients* (24, 48). When dealing with spiking signals, a naive attempt to minimize the mean-squared error 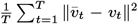 with gradient-based methods leads to exploding cost function gradients as the dataset horizon *T* increases. This occurs due to the high sensitivity of spiking models to changes in their parameters (47), which in turn is due to the existence of bifurcations in neural dynamics (31). The two methods outlined below adapt the techniques of teacher forcing (22) and multiple-shooting (23) to the context of RMMs, eliminating exploding gradients altogether.

### Teacher forcing (TF) with backpropagation

Teacher forcing (TF) (22) adapted to RMMs is formulated as the supervised learning problem

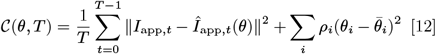

where *I*_app,*t*_ is the measured applied current, *Î*_app,*t*_ is an applied current estimate, *ρ*_*i*_ are regularization constants, and 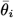 are parameter priors. We impose *ρ*_*i*_ = 0 whenever *θ*_*i*_ is either a layer bias or an element of the 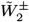 matrices of a mechanistic current. We also impose 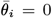 whenever *θ*_*i*_ is not a Nernst potential. TF, which has been applied to conductance-based models in (20), uses voltage measurements to estimate the applied current. Manipulating Eq. (2), we find that the current estimate is

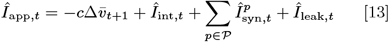

where

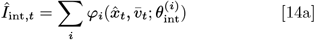

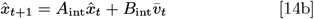

is used to compute the internal current estimate, with 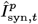 and *Î*_leak,*t*_ being computed analogously using the measured 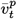 and 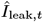, respectively. Eq. (12) implements TF because training is carried out by replacing *v*_*t*_ by the measured (teacher) signal 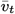 in the model dynamics. The formulation has roots in the *equation-error* method of system identification (49).

In the SI Appendix we show that when applied to RMMs with fixed *A* and *B* matrices, TF allows massively parallelized GPU computations during training, which is the reason why RMMs can be trained so quickly. In short, when *A* and *B* need not be trained, the estimated applied current *Î*_app,*t*_ in Eq. (13) effectively becomes a feed-forward neural network, whose inputs are given by the the voltage signal 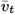 and the corresponding pre-simulated internal states. Since those inputs can be concatenated into a large matrix, this implies that Eq. (12) can be solved using one’s favourite backpropagation-based gradient descent algorithm, while exploiting efficient matrix-based computations in the GPU.

#### Multiple shooting with backpropagation through time

Teacher forcing (TF) comes with the downside that since the model is trained to predict the applied current, it might not predict voltage satisfactorily. Although in our results we have shown that TF yields very good results on experimental data, better models can be found by exploiting multiple shooting (23), which we adapt to RMMs, and which generalizes the TF method above. In multiple shooting, a dataset of length *T* is divided into *N* smaller intervals of length *T*_*s*_, with *N* the largest integer such that *NT*_*s*_ ≤ *T*. To construct a training cost function, the model being fit is simulated *N* times, with the initial conditions of each simulation defined independently. Mathematically, for each *n* = 0, 1, …, *N −* 1 and each *t* = 0, 1, …, *T*_*s*_ *−* 1, we define the inputs 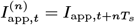 and simulate the system to obtain *N* state vectors, which we denote by

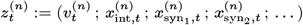

and which are initialized according to 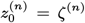, where the *ζ*_(*n*)_ are arbitrary initial conditions. In practice, good initial conditions are given by the measurements and the teacher-forced states, i.e.

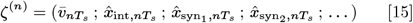

with 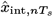 obtained from Eq. (14).

The key feature of multiple shooting is that the *ζ*^(*n*)^ are trained alongside *θ* so as to minimize the discrepancy between the state at the end of the *n*^th^ simulation, 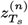, and the initial condition of the (*n* + 1)^th^ simulation, *ζ*^(*n*+1)^. This is accomplished by solving the optimization problem given by

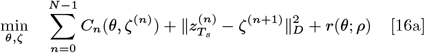

where 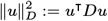 is a *D*-weighted 2-norm,

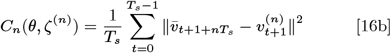

are the cost functions associated to each of the *N* simulations, and *r*(*θ*; *ρ*) is the same parameter regularization term as in Eq. (12). The problem Eq. (16) is an unconstrained optimization problem on the parameters *θ* and initial conditions *ζ*^(*n*)^; it can be solved efficiently with gradient descent. When training with multiple shooting, computing the gradient of the cost function in Eq. (16a) requires performing backpropagation through time over the *T*_*s*_ timesteps of each parallel simulation; to avoid exploding gradients one must keep *T*_*s*_ small enough. The size of the shot size *T*_*s*_ is a hyperparameter of the problem; larger shot sizes may yield better predictions at the cost of slower learning^3^. Multiple shooting can be sped up by exploiting parallelization, as explained in the SI Appendix.

### Validating RMMs

All models in this paper were validated on held-out training data as follows: a “snapshot” of models during training was saved so that each training run yielded around 100 snapshot models; after training, snapshot models were simulated using Eq. (2) and the validation *I*_app_ (this was done using parallel computing to speed up the validation process), and the best model was chosen as the result of the training procedure. This was done as a means to prevent overfitting, and to recover loss curves. Validation of all 100 snapshot models did not exceed two minutes for any of the results in this paper. The best snapshot model was determined by comparing its predicted spike times (including intra-burst spikes) to the validation voltage spike times. This was done by computing a validation cost given by

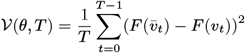

where *v*_*t*_ is the predicted voltage trace, 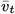 is the target validation voltage trace, and *F* (·) is a nonlinear filter designed to extract a smoothed spike train out of the voltage traces. The nonlinear filter *F* applies, in succession: a bandpass 6^th^ order Butterworth filter with cutoff frequencies at 1*/*50 and 1*/*2 kHz (which filters out most of the slow components of the voltage traces); a relu nonlinearity with threshold set at 2 mV (which transforms the filtered spike train into a sequence of approximate impulse functions); and a Gaussian smoother with standard deviation of 50 (HH neuron RMM), 200 (LP neuron RMM) and 500 mV (PD neuron RMM) which smooths the approximate impulse train. Notice comparing smoothed spike trains is an established methodology to validate statistical integrate-and-fire models (12).

#### Hogkin-Huxley RMM

In Figure 2, training data was generated by simulating the Hodgkin-Huxley model (26, 31) with the forward-euler discretization scheme and simulation timestep of *δ*_sim_ = 0.01 ms. A diagonal state space realization was constructed with the time constants mentioned in Results; mechanistic readouts are given by Eq. (5) and Eq. (9), with a single output and 20 ANN units; Nernst potentials were fixed to those of the HH model. The training *I*_app_ consisted in three realizations of a stochastic process defined by

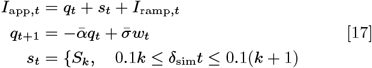

where *I*_ramp,*t*_ is a deterministic ramp of applied current such that *I*_ramp,0_ = *−*15*µA* and *I*_ramp,*T*_ = 0*µA*; *q*_*t*_ is the output of a discretized Ornstein-Uhlenbeck process as described above, with 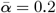, and 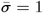; and *s*_*t*_ is a random process given by a sequence of steps of random uniformly distributed amplitude *S*_*k*_ ∼ *𝒰* [0, 10]. For learning, the resulting simulated data was downsampled to *δ* = 0.05 ms, and the voltage was corrupted by white Gaussian noise of standard deviation equal to 0.5 mV. One realisation of the training dataset is illustrated in Figure S1. The validation dataset was obtained by stimulating the HH model with Eq. (17) containing only the Ornstein-Uhlenbeck component, with 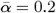 and 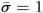 (*I*_ramp,*t*_ = 0 and *s*_*t*_ = 0). The Hodgkin-Huxley RMM was trained with TF and a regularization constant of *ρ*_*i*_ = 10^*−*6^ for all weights in the model not belonging to 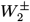. The cost Eq. (12) was minimized with the ADAM routine with step size 0.001 and momentum parameters *β*_1_ = 0.9, *β*_2_ = 0.999. Training curves can be found in the SI Appendix.

### LP neuron RMM

To construct the LP neuron RMM, the intrinsic and synaptic currents were defined based on the sequence of continuous time constants 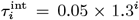, *i* = 0, 1, …, 35, and *τ*^syn^ = 0.05 × 1.3^*i*^, *i* = 14, 15,…, 29,, which were chosen pragmatically so as to ensure that all timescales usually found in LP model are represented; see Figure S4 for representative time constant functions found in an LP conductance-based model (42). A state space realization with orthogonal impulse responses was used. The model is constructed with two lumped readouts corresponding to the total intrinsic and total synaptic currents, with the corresponding MLPs given by Eq. (8). The MLPs were chosen with *L*_int_ = 3 layers and *L*_syn_ = 2 layers, respectively, with ten units in any given layer. In Figure 3, training and validation data for the LP neuron consisted in voltage 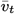 and applied current *I*_app,*t*_ obtained experimentally (see *electrophysiology* in Methods). The applied current used to train and validate the RMM was given by a discretized OU process, that is *I*_app,*t*_ = *q*_*t*_, with *q*_*t*_ as in Eq. (17), with 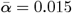 and 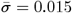. To train the model, we first used TF with *ρ*_*i*_ = 0.0015 for all weights in the model, and minimized the cost function with the ADAM gradient descent routine with step size 0.003, *β*_1_ = *β*_2_ = 0.9, and a mini-batch size of 484000 datapoints. In Figure 3 B we shifted the synaptic current prediction so that its minimum value is at zero. In Figure 3 we used multiple shooting with *T*_*s*_ = 30 and the same *ρ*_*i*_, mini-batch size and and ADAM hyperparameters as TF; final states were regularized with 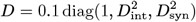 where *D*_int_ and *D*_syn_ are diagonal matrices whose diagonals are given by the inverse DC gains [(*I − A*)^*−*1^*B*]^*−*1^.

### PD neuron RMM

To construct the PD neuron RMM, the intrinsic current matrices *A* and *B* were defined based on the sequence of continuous time constants 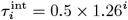, *i* = 0, 1, …, 31 (this results in logarithmically spread out time constants from 0.5 ms to around 643 ms). This was again chosen pragmatically based on the range of time constants from (42) (see Figure S4). The PD neuron RMM contained four currents: a lumped current *I*_lumped_, given by Eq. (4), and three mechanistic data-driven currents *I*_K_, *I*_Ca_, and *I*_H_, given by Eq. (11). To provide states for the current readouts, a block-diagonal *A* matrix was constructed using a mixed realization with 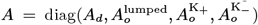, where *A*_*d*_ is a diagonal realization constructed with 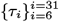, and 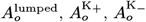 are realizations with orthogonal impulse responses constructed with 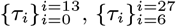, and 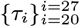, respectively. The readouts of *I*_Ca_ and *I*_H_ read from the diagonal part of the state space system, while the readouts of *I*_lumped_ and *I*_K_ read from the part with orthogonal impulse responses. Each of the activation (*σ*_+_) and inactivation (*σ*_*−*_) mappings in the mechanistic currents contained two layers, with 20 and 10 units in each layer, respectively; the *H* current had no activation mapping. Conductance scaling parameters were initialized at 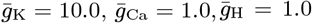 and were left fixed throughout training. Nernst potentials of the mechanistic currents were initialized to *E*_K_ = − 80, *E*_K_ = 130 + *ϵ*, and *E*_H_ = −10 + *ϵ*, with *ϵ* a vector of zero-mean normally distributed random variables with standard deviation of 10 mV. Nernst potential vectors of the *I*_H_ and *I*_Ca_ currents were allowed to be trained, while the Nernst potential of *I*_K_ was fixed. In the mechanistic currents, the normalization layers *h*_±_ (·) at the input of *σ*_±_ (·) were defined with biophysical priors for the voltage range given by 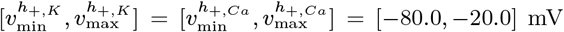, and 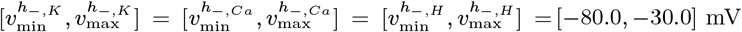 (this choice was based on the activation functions of the model from (42)).

In Figure 4, training and validation data for the AB-PD subsystem consisted in voltage 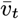 and applied current *I*_app,*t*_ obtained experimentally from a single PD neuron following the experimental procotol described in the *electrophysiology* section. The dataset used to train and validate the RMM consisted of a combination of four different 42-second long trials where the PD neuron was excited with the discretized OU process Eq. (17); in two of the trials 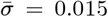 was fixed and 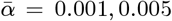 was varied, and in two of the trials 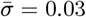 was fixed and 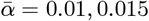 was varied. To train the model, we used TF with *ρ*_*i*_ = 10^*−*5^ for all weights belonging to hidden ANN layers; for weights in 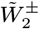 we used *ρ*_*i*_ = 0, while for weights in 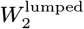 we used *ρ*_*i*_ = 0.01. The cost was minimized with the ADAM gradient descent routine with step size 0.001, momentum parameters of *β*_1_ = 0.9, *β*_2_ = 0.99, and a mini-batch size of 484000 datapoints.

### Electrophysiology

#### Animals

Adult male Jonah Crabs, *Cancer borealis*, were acquired from Commercial Lobster (Boston, MA) between August 2022 and July 2023. Animals were held in groups up to 5 in 75 Gallon tanks filled with artificial seawater at 11-13°C with a 12 hr light/dark cycle. The crabs were put on ice for at least 30 min before the dissection. Dissections were performed as described in (50). In brief, the crab stomach was removed, the stomatogastric nervous system (STNS) was dissected from the stomach and pinned down in a petri dish (10 ml) lined with Sylgard (The Dow Chemical Company). The STNS includes the stomatogastric ganglion (STG) containing the cells of interest, modulatory inputs descending into the STG from the esophageal ganglion and the two bilateral commissural ganglia, and the motor nerves coming from STG. The STNS was held at 4°C overnight in solution (saline) containing 440 mM NaCl, 11 mM KCl, 26 mM MgCl_2_, 13 mM CaCl_2_, 11.2 mM Trizma base, 5.1 mM maleic acid, pH 7.45, and used for the experiment next day.

#### Recordings

The STG was desheathed and intracellular recordings from somata of the lateral pyloric (LP) and pyloric dilator (PD) neurons were obtained. Sharp glass microelectrodes (10-20 MΩ) filled with 0.6 M K_2_SO_4_ and 20 mM KCl were used to record the intracellular activity. During the experiment the STNS-containing dish was pump-perfused with 11°C saline at a rate of 5ml/min. The temperature was controlled by Peltier device connected to the model CL-100 temperature controller (Warner Instruments). The cells were identified by their activity (monitored by extracellular electrodes) on the motor nerves: PD activity on the pyloric dilator nerve and LP activity on the lateral ventricular nerve. Extracellular recordings were done with a stainless-steel electrode in a well (Vaseline and 10% mineral oil mixture) around the motor nerve of interest. An Axoclamp 900A amplifier (Molecular Devices, San Jose) was used to amplify intracellular signals. Extracellular signals were amplified by an A-M Systems 1700 extracellular amplifiers. Data were obtained with pClamp data acquisition software (Molecular Devices, San Jose, version 10.5), Digidata 1440A digitizer (Molecular Devices, San Jose) and Real-Time eXperiment Interface (RTXI) software version 2.2. Recordings were acquired at 10 kHz sampling frequency. A custom RTXI module was used for the Ornstein-Uhlenbeck noise current injection. After the cells were identified, the solution was switched to saline containing 10^*−*5^M picrotoxin (PTX), Sigma-Aldrich, to block the inhibitory synaptic connections between cells. Recording started 30 min after PTX was applied. The membrane voltage was recorded (in PD and LP) and current was injected (in LP or in PD, depending on the experiment) with a single electrode in Discontinuous Current Clamp (DCC) mode. To filter out the inherent noise of DCC recordings, PD neuron data was filtered with a 4^th^ order Butterworth filter with cutoff frequency given *f*_*c*_ = 4 kHz.

#### Hardware

Training times in this paper were obtained with a desktop computer set up with an AMD Ryzen 7 7800×3D 8-Core Processor, 64 Gigabites of RAM, and a NVIDIA GeForce RTX 4090 graphics card with 24 GB of VRAM.

## Acknowledgements

The research leading to these results has been funded by the Kavli Foundation and the National Institutes of Health (R35 NS 097343). Their support is greatly appreciated. The authors thank Rodolphe Sepulchre, Fulvio Forni, Dhruva Raman, Alessio Franci and Kyra Schapiro for their insightful comments on this work.

## Supporting Information for

**This PDF file includes:**

Supporting text

Figs. S1 to S6

SI References

## Supporting Information Text

### Methods

#### Weight initialization

In lumped and mechanistic currents, the purpose of weight initialization is to obtain a random set of ANN weights and biases which is empirically optimized for the ANN to learn efficiently from inputs that lie mostly on the interval [*−*1, +1]. Initialization for lumped and mechanistic currents differ due to the weight constraints mentioned in the main text.

#### Lumped currents

The *W* matrices in each layer of the lumped readouts are initialized with the scaled Nguyen-Widrow heuristic (1). Letting *W* [*k*] denote the weight matrix at iteration *k* of the learning algorithm, at each layer *𝓁* we set

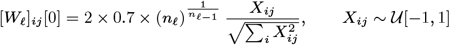

where *n*_*𝓁*_ is the number of units in the layer *𝓁* (the number of inputs of the MLP when *𝓁* = 0). The 2*×* factor is included in the heuristic to optimize the weights for use with tanh, whose effective activation range is the interval [*−*2, 2]. The biases *b* in each layer are spread evenly according to

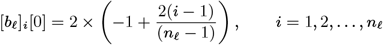

if *n*_*𝓁*_ *>* 1, and [*b*_*𝓁*_]_1_ = 0 if *n*_*𝓁*_ = 1.

#### Mechanistic currents

The weight initialization heuristic changes in mechanistic currents due to the elementwise positivity constraints on the weight matrices. The weight matrices 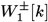 and 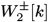 at iteration *k* of the learning algorithm are given by

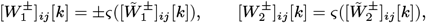

where *ς*(·) = log(1 + exp(·)) is the softplus function. The matrices 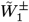 and 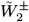 contain the parameters being learned.

We initialize 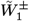 according to

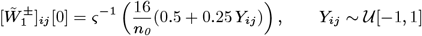

where *n*_*0*_ is the number of outputs of the mechanistic normalization layer (inputs of *σ*_+_), and *ς*^*−*1^(·) = · + log(1 − exp(−·) is the inverse of the softplus function. The corresponding biases are given by

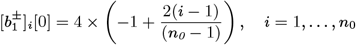

The initialization heuristic above ensures that

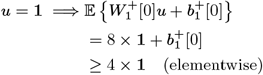

which implies that at initialization, 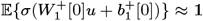 whenever *u* = **1**, since the effective activation range of the logistic nonlinearity *σ*(*·*) is the interval [*−*4, 4]. Similarly, we have

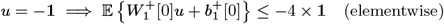

and thus 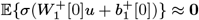 whenever *u* = *−***1**. A completely analogous consideration applies to 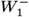.

We initialize 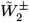 according to

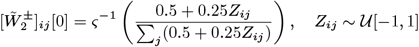

which implies that at initialization, 𝔼 {*σ*_+_(*u*)} ≈ **1** whenever *u* = **1** and 𝔼 {*σ*_+_(*u*)} ≈ **0** whenever *u* = − **1**. An analogous reasoning applies to 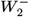 and *σ*_−_ (·).

The above initialization heuristic guarantees that mechanistic currents are initially shut down for steady-states corresponding to voltages at the boundaries of a prior activation range; see *Normalization layers* below.

#### Normalization layers

The normalization layers of lumped currents and mechanistic currents differ in the interval of readout inputs which is mapped to [*−*1, +1].

#### Lumped currents

In lumped currents, the fixed mappings *h*(*v*_*t*_, *x*_*t*_) are given by

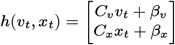

where *C*_*v*_ and *β*_*v*_ are used to perform min-max normalization of *v*_*t*_, and *C*_*x*_ and *β*_*x*_ are used to select a subset of states of *x*_*t*_ and perform min-max normalization on those states. Min-max normalization is accomplished as follows: given the data 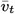 used for training, we first obtain a state trajectory 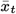 by simulating Eq. 3b (main text) with the voltage data 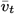 (transients not used during training are discarded). Then the *C* matrices and *β* shift vectors are calculated to ensure that the interval 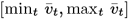, as well as each of the intervals 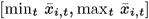, are mapped onto [*−*1, 1]. Mathematically, we have

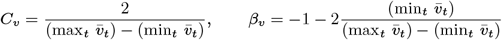

for voltage, and

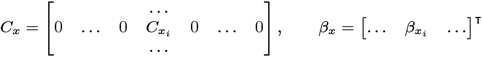

with 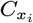 and 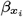 computed analogously to *C*_*v*_ and *β*_*v*_ above (here, *i* is the index of a state selected to be an input to the MLP).

#### Mechanistic currents

In mechanistic currents, for each activation or inactivation readout, *σ*_+_(*h*_+_(*x*_*t*_)) and *σ*_*−*_(*h*_*−*_(*x*_*t*_)) respectively, the fixed mapping

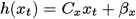

selects a subset of states of *x*_*t*_ and normalizes them. However, instead of min-max normalization, we employ normalization based on a prior voltage interval over which we initially believe a current activates or inactivates. Given such an interval [*v*_low_, *v*_high_], we first compute the steady-states

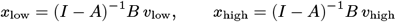

at the endpoints of [*v*_low_, *v*_high_]. Then, *C*_*x*_ and *β*_*x*_ are found by mapping each of the intervals [*x*_*i*,low_, *x*_*i*,high_] onto [−1, 1], where *i* is the index of a state selected to be an input to *σ*(·). Mathematically, the expressions for *C*_*x*_ and *β*_*x*_ for mechanistic currents are analogous to those of lumped currents.

Notice that because we employ state space realisations with elementwise positive steady-state gain (*I*− *A*)^*−*1^*B*, the C matrices found in this way are also elementwise positive.

#### Monotonicity of mechanistic currents with diagonal state space realisations

Consider the function *σ*_+_(*h*_+_(*x*)) modelling the activation of a mechanistic readout which reads from a diagonal state space realisation with *A* = diag {*λ*_*j*_} and *B* = (**1**− vec(*λ*_*j*_)). The input-output system realised by 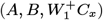 is said to be *positive* (2), because we are dealing with a discrete time system whose state space matrices are all elementwise positive. As a consequence, monotonically increasing/decreasing input (voltage) trajectories yield monotonically increasing/decreasing trajectories at the input of the logistic function nonlinearities The elementwise positivity of 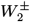 coupled with the monotonicity of the logistic nonlinearities in *σ*_+_(*h*_+_(*x*)) then ensures that monotonicity of those trajectories is preserved at the output of the readout. The argument is analogous for *σ*_*−*_(*h*_*−*_(*x*)), with the difference that since 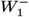 is elementwise negative, the realisation 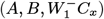 is a *negative system*. In that case monotonically increasing/decreasing voltage trajectories yield monotonically decreasing/increasing trajectories at the input of the logistic nonlinearities; this is again preserved by the monotonicity of the logistic function and elementwise positivity of 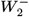.

### State space system with orthogonal impulse responses

The discrete-time poles *λ*_*j*_ are can be used to *A* and *B* so as to realize a linear system with orthogonal impulse responses (also known as convolution kernels), pointed out in the main text. To construct a system with no timescale separation between states, this is accomplished by defining *A* = *A*_*n*_ and *B* = *B*_*n*_, where *A*_*n*_ and *B*_*n*_ are given by the iterative procedure

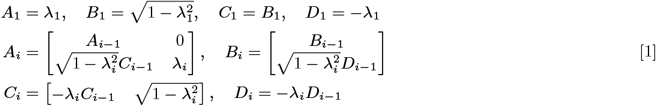

for *i* = 1, …, *n* (see (4, Chapter 2) for details). The above procedure results in a system whose state components (the elements of *x*_*t*_) follow distinctly different trajectories given sudden changes in the input; we have found empirically that this results in significantly better predictive models when compared to simpler choices of *A* and *B*. Figure S4 of the SI Appendix shows how the responses of a system with orthogonal impulse responses overlap less in time, providing ANNs with a richer set of input signals. Notice that for slow state components, a spike is well approximated by the impulse function, and hence spike responses of slow states will be near orthogonal (modulo a constant dependent on the mean of the spiking signal).

Whenever timescale separation between components of the state vector *x*_*t*_ is desired, the procedure is slightly modified. To construct *A* and *B* in those cases, we first partition the time constant sequence 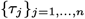 into non-intersecting subsequences 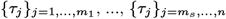 where *s >* 0 is the number of desired timescale partitions. Then, we define *A* as a block-diagonal matrix whose diagonal (matrix) elements are each constructed according to Eq. (1) using the corresponding subsequences of discrete-time poles 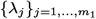, etc. This modification ensures that the state vector partitions 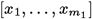, etc., do not share dynamical modes in the same timescales.

#### Parallelization with Teacher forcing

Teacher forcing lends itself exceptionally well for the purposes of training RMMs, and in case the state space matrices *A* and *B* remain fixed (non-trainable), massively parallelized computations in GPUs can be exploited to their fullest potential. To see why, assume *A* and *B* are fixed a priori, and let

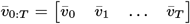

denote a row vector containing all voltage measurements. Teacher-forced state estimates can be obtained from

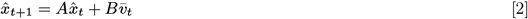

which need to be computed only once as *A* and *B* will not be trained. The state estimates can also be put in matrix form as

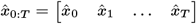

with 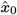 a “warmed-up” initial condition^*^. By precomputing 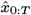, minimizing the teacher forcing cost function (Eq. 12 in main text) in *θ* reduces to the problem of training a feed-forward ANN with inputs and targets given by large matrices, enabling massive parallelization. Indeed, Eq. 12 (main text), and Eq. 13 (main text) can be rewritten as

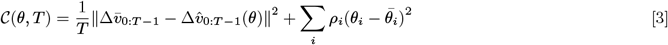

with 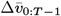 a row vector of target ANN outputs, and 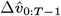 being the output of a feed-forward ANN with matrix inputs 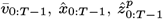, and *I*_app,0:*T −*1_. Notice that in practice, the capacitance *c* is lumped into the output layers of the *φ* sub-networks, and recovered as the inverse of the ANN input weight corresponding to *I*_app_.

Computing the gradient of Eq. (3) can be done efficiently with backpropagation, which for moderate number of layers in the model’s MLPs causes no exploding nor vanishing gradients^†^. Notice that this “static” version of teacher forcing relies on two facts: first, the fact that bounds on physiological time constants are well-known, which allow fixing *A* and *B* in a reasonable manner; and second, the fact that *A* can be trivially constrained to be stable (Hurwitz), a property that ensures that the states 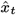 and 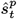 remain bounded for any possible measurements 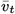 and 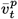.

#### Parallelization with Multiple shooting

Letting

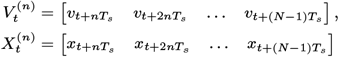

and similarly defining *S*_*t*_ and 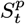, the model can be simulated in parallel on each of the *N* multiple shooting intervals by simulating Eqs.1-6 (main text) blockwise, that is, replacing *v*_*t*_ by 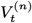, *x*_*t*_ by 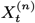, and so on. It follows that in multiple shooting there is a natural tradeoff between the speed of training (enabled by large *N* and massive parallelization) and the quality of predictions, which tends to increase (up to a bound determined by exploding gradients) with longer intervals *T*_*s*_ (see (5), (6) for a formal analysis).

**Fig. S1.**
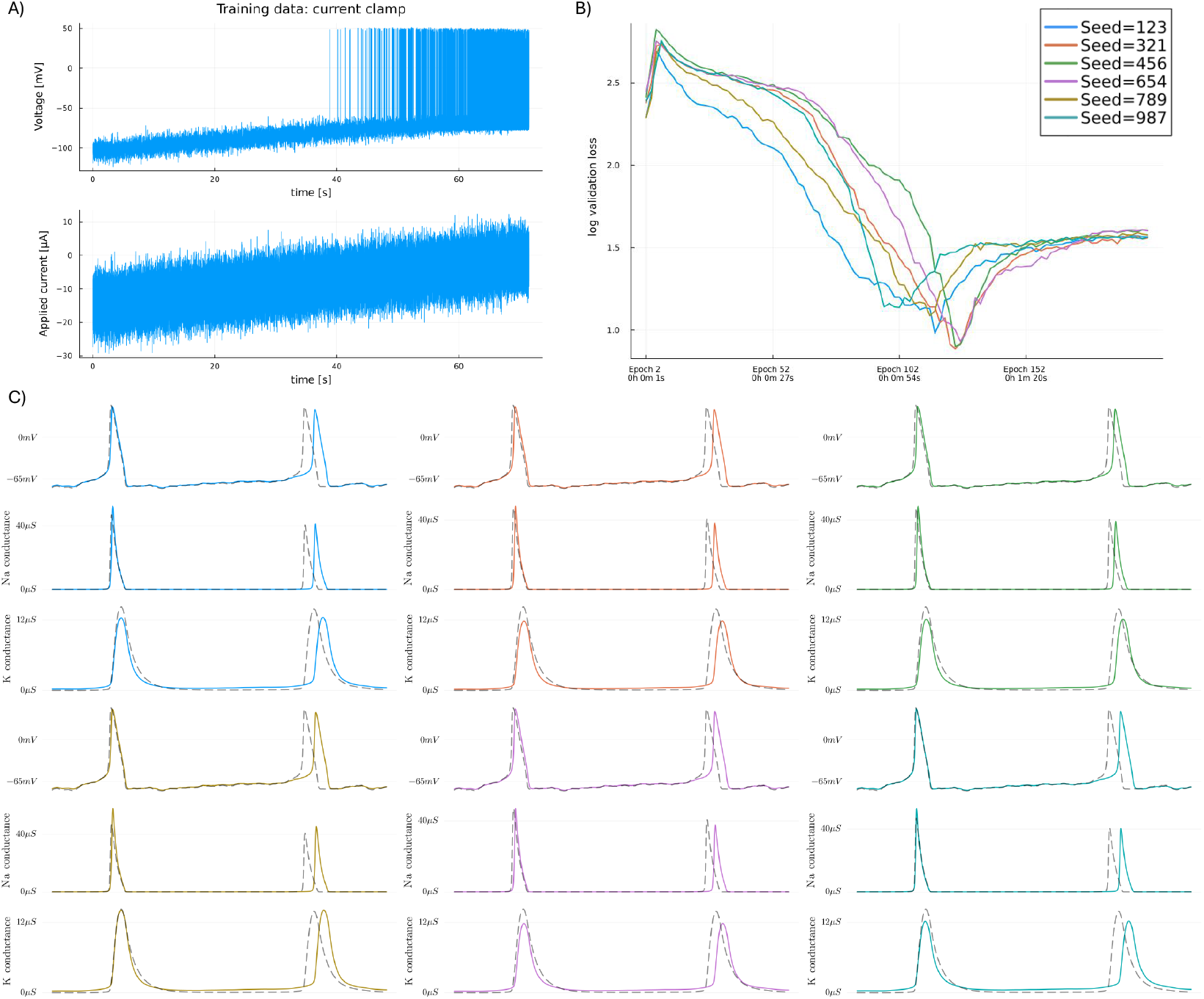
A) One realization of the input-output data used to train the Hodgkin-Huxley RMM. Three realizations of the same stochastic applied current were used to train the model. B) Validation loss of six Hodgkin-Huxley RMMs trained with the data of (A). Training used the same hyperparameters, but different random number generator (RNG) seeds. RNG seeds were used to randomize parameter initialization, as well as to randomize the shuffling algorithm of stochastic gradient descent. C) Comparison of the predictive accuracy of the voltage and conductance traces of all six models whose validation curves are plotted in (B). Models were selected at the minimum of the validation curves.

**Fig. S2.**
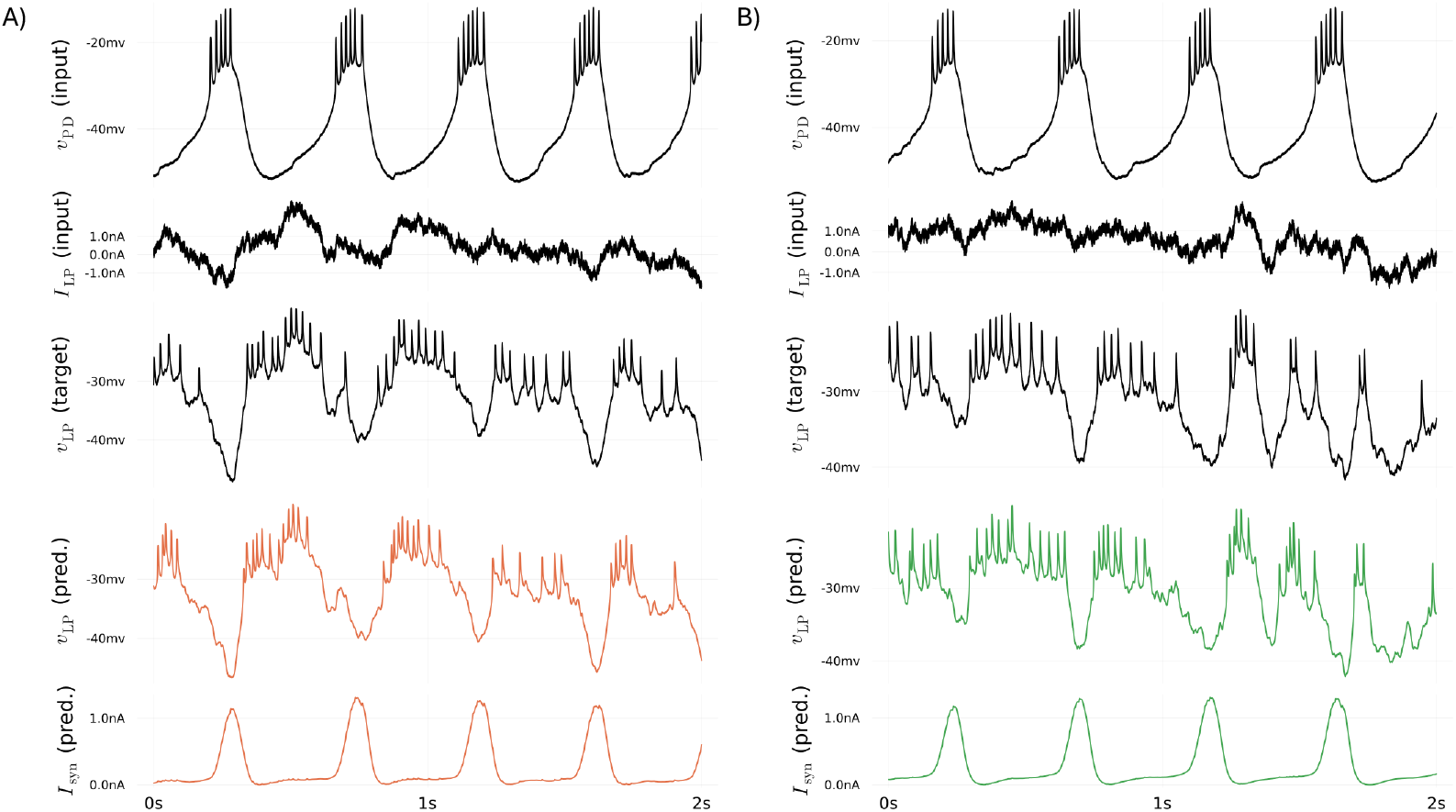
Prediction results for the two LP models whose training curves are presented in the main body of the text.

**Fig. S3.**
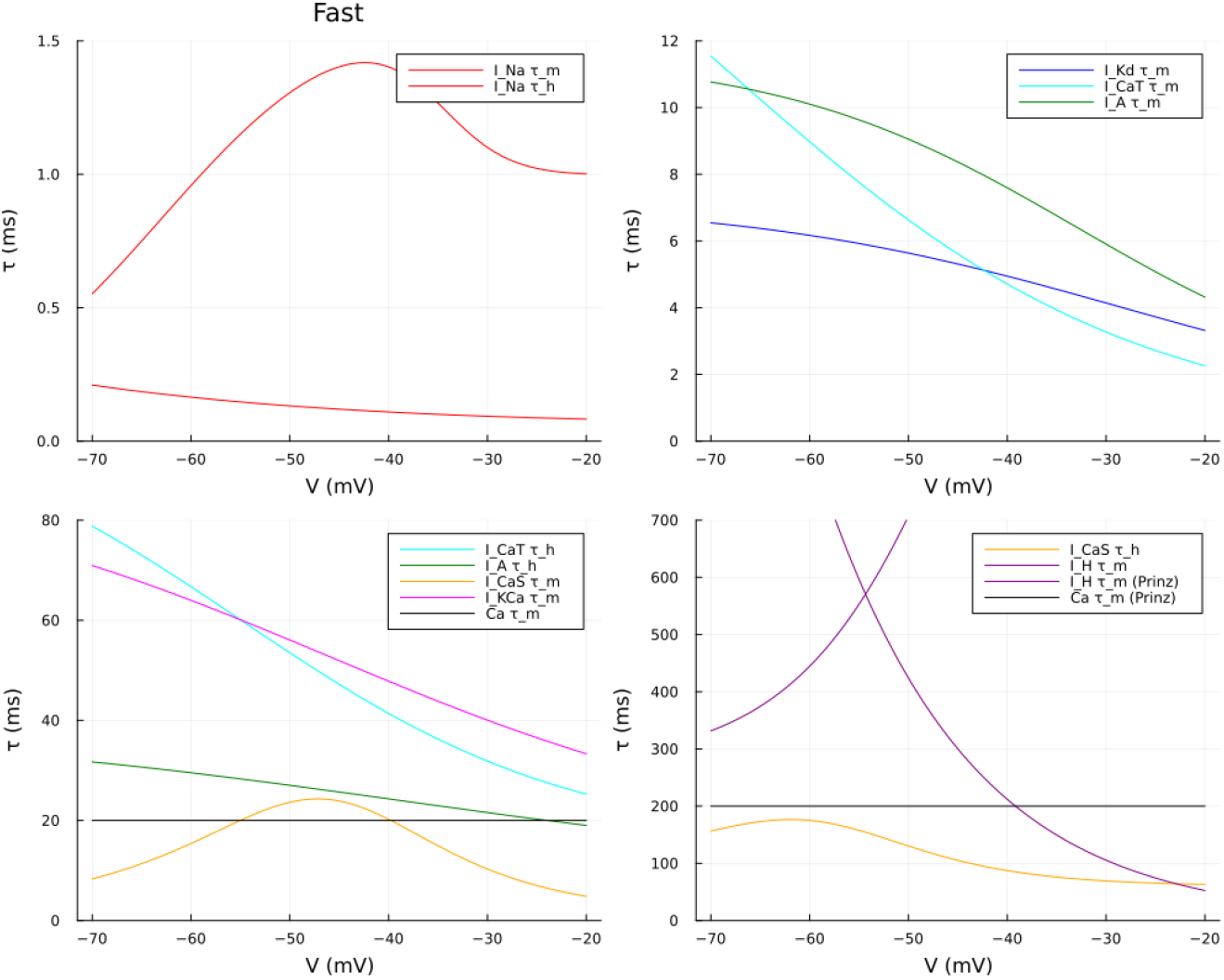
Time constant functions of the conductance-based model of (7). These functions are also used in (8), who considered different time constants for calcium concentration and the H current, and doubled the values of all time constant functions plotted above.

**Fig. S4.**
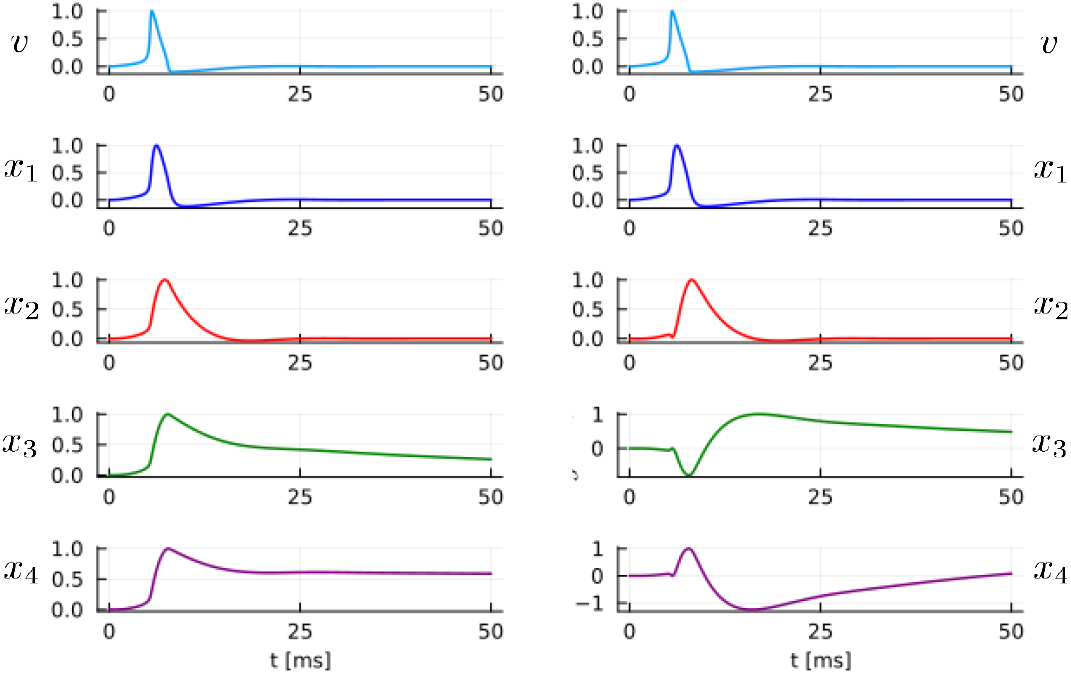
Spike responses of two different choices of state space systems. **Left:** state component responses *x*_*i,t*_ of a four-dimensional diagonal state space system given by *A* = diag(*λ*_*i*_), *B* = ((1− *λ*_1_), …, (1− *λ*_*n*_))^⊺^). Here, *λ*_*i*_ = exp(−*δ/τ*_*i*_) is computed using a sampling period of *δ* = 0.01 and continuous time constants *τ*_1_ = 0.5, *τ*_2_ = 5, *τ*_3_ = 50, and *τ*_4_ = 500. Spike responses, which are shown normalized by maximum value, are the output of the system to the spike shown on top. **Right:** spike responses of orthogonal filters, constructed with the same sequence of *λ*_*i*_ according to the procedure described in Methods. In the immediate aftermath of a spike, orthogonal filters provide richer signal features for artificial neural networks to be trained and recombine into ionic currents.

**Fig. S5.**
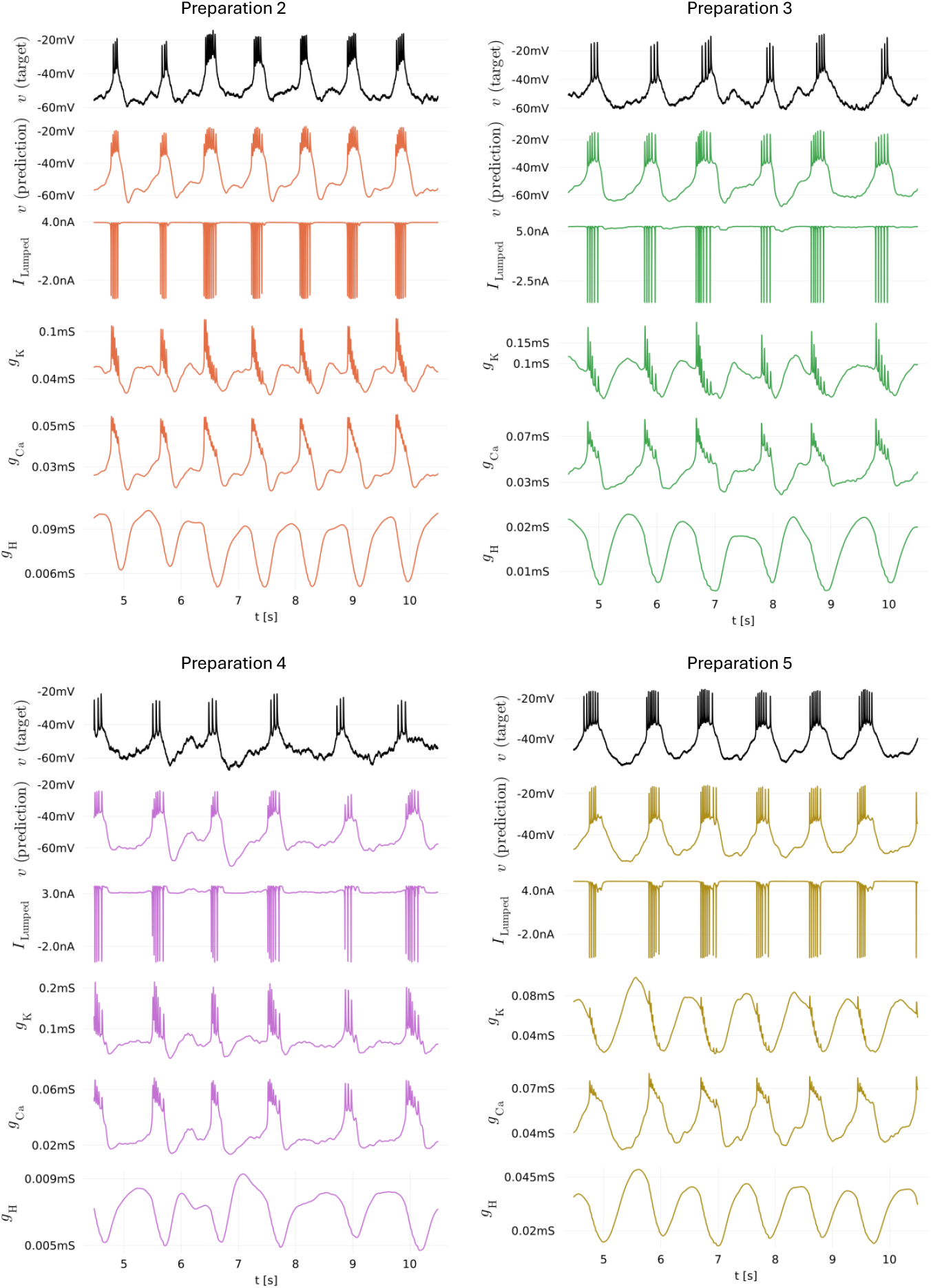
Prediction results for four different PD model preparations in addition to the preparation whose predictions are shown in Figure 3 of the main text. The RMMs were trained on these preparations using the same hyperparameters used to obtain the results of Figure 3, except for the overall regularization constant (*ρ*_prep2_ = 7.5 *×* 10^*−*5^, *ρ*_prep3_ = 5 *×* 10^*−*5^, *ρ*_prep4_ = 5 *×* 10^*−*5^,*ρ*_prep5_ = 2 *×* 10^*−*5^). While electrophysiological properties of the PD neurons are visibly different, the RMM is able to capture their dynamics reasonably well. All models whose predictions are shown in this figure have been trained in around 2 minutes (see Hardware in Methods, in the main text).

**Fig. S6.**
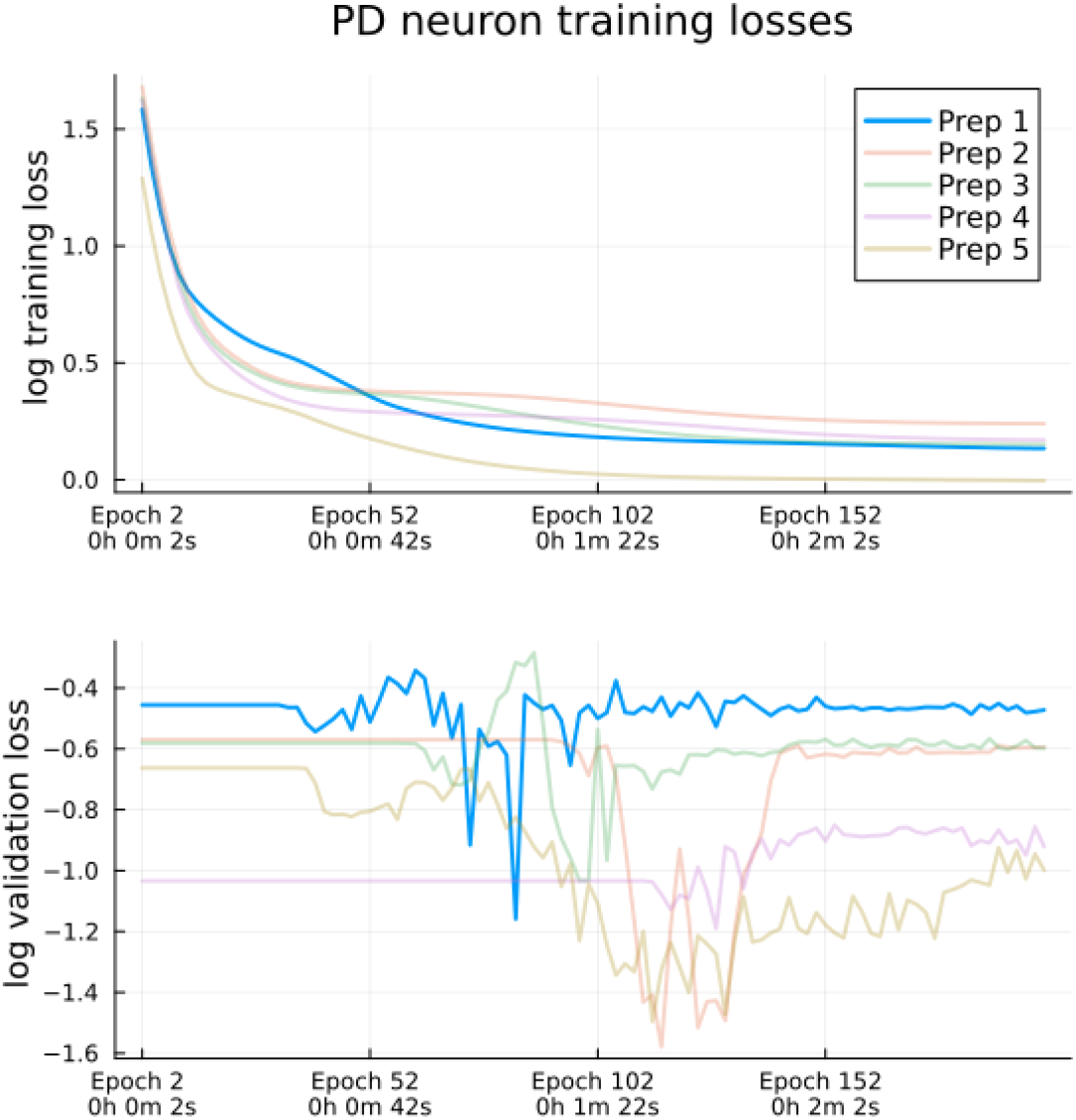
Training and validation losses of the five PD neuron RMMs trained on experimental data. Blue traces refer to the RMM shown in Figure 3 of the main text, while other traces refer to RMMs shown in Figure S5.

## Footnotes

1 When such a model is not available, time constants can alternatively be based on the frequency content of the voltage recording

2 The impulse response of a LTI system is the kernel of that system’s convolution representation.

3 It can be seen in Eq. (16) that when initial conditions are chosen according to Eq. (15), multiple shooting generalizes TF, which is recovered by choosing *N* = *T* and *T*_*s*_ = 1.

* Warming up consists in choosing 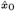 by simulating the state space model with a reserved part of the voltage data: suppose the initial segment of the dataset is given by 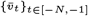 then this simply means computing Eq. (2) for *t* = *−N*, …, *−*1.

† In practice, additional tricks used to train feed-forward neural networks with gradient descent can be employed to improve generalization of the trained model. Two of those tricks are *mini-batching*, which divides the dataset into multiple batches so as to perform more frequent descent steps, and *shuffling*, which randomly permutes the columns of input and output matrices at the beginning of each training epoch. The joint application of such tricks has come to be known as *stochastic gradient descent*.

## References

1. L Paninski, J Pillow, E Simoncelli, Comparing integrate-and-fire models estimated using intracellular and extracellular data. Neurocomputing 65, 379–385 (2005).

2. H Abarbanel, D Creveling, R Farsian, M Kostuk, Dynamical State and Parameter Estimation. SIAM J. on Appl. Dyn. Syst. 8, 1341–1381 (2009).

3. CD Meliza, et al., Estimating parameters and predicting membrane voltages with conductance-based neuron models. Biol. Cybern. 108, 495–516 (2014).

4. JM Lueckmann, et al., Flexible statistical inference for mechanistic models of neural dynamics in 31st Conference on Neural Information Processing Systems (NIPS 2017). (Long Beach, CA, USA), p. 18 (2017).

5. K Abu-Hassan, et al., Optimal solid state neurons. Nat. Commun. 10, 5309 (2019).

6. LM Alonso, E Marder, Visualization of currents in neural models with similar behavior and different conductance densities. eLife 8, e42722 (2019).

7. JD Taylor, S Winnall, A Nogaret, Estimation of neuron parameters from imperfect observations. PLOS Comput. Biol. 16, e1008053 (2020).

8. PJ Gonçalves, et al., Training deep neural density estimators to identify mechanistic models of neural dynamics. eLife 9, e56261 (2020).

9. TB Burghi, R Sepulchre, Adaptive observers for biophysical neuronal circuits. IEEE Transactions on Autom. Control. 69, 5020–5033 (2024).

10. SA Wells, JD Taylor, PG Morris, A Nogaret, Inferring the Dynamics of Ionic Currents from Recursive Piecewise Data Assimilation of Approximate Neuron Models. PRX Life 2, 023007 (2024).

11. J Pillow, M Park, Adaptive Bayesian Methods for Closed-Loop Neurophysiology in Closed Loop Neuroscience. (Academic Press, Cambridge, MA), pp. 3–18 (2016).

12. W Gerstner, WM Kistler, R Naud, L Paninski, Neuronal Dynamics: From Single Neurons to Networks and Models of Cognition. (Cambridge University Press, Cambridge, UK), (2014).

13. TB Burghi, M Schoukens, R Sepulchre, System identification of biophysical neuronal models in 59th IEEE Conference on Decision and Control. (Jeju Island, South Korea), pp. 6180–6185 (2020).

14. D Beniaguev, I Segev, M London, Single cortical neurons as deep artificial neural networks. Neuron 109, 2727–2739.e3 (2021).

15. KW Latimer, F Rieke, JW Pillow, Inferring synaptic inputs from spikes with a conductance-based neural encoding model. eLife 8, e47012 (2019).

16. D Durstewitz, G Koppe, MI Thurm, Reconstructing computational system dynamics from neural data with recurrent neural networks. Nat. Rev. Neurosci. 24, 693–710 (2023).

17. M Aguiar, A Das, KH Johansson, Learning flow functions of spiking systems in Proceedings of the 6th Annual Learning for Dynamics & Control Conference. (PMLR), pp. 591–602 (2024).

18. M Brenner, et al., Tractable Dendritic RNNs for Reconstructing Nonlinear Dynamical Systems in Proceedings of the 39th International Conference on Machine Learning. (PMLR), pp. 2292–2320 (2022).

19. R Naud, W Gerstner, Can we predict every spike? in Spike Timing: Mechanisms and Function. (CRC Press), (2013).

20. QJM Huys, MB Ahrens, L Paninski, Efficient Estimation of Detailed Single-Neuron Models. J. Neurophysiol. 96, 872–890 (2006).

21. AA Prinz, D Bucher, E Marder, Similar network activity from disparate circuit parameters. Nat. Neurosci. 7, 1345–1352 (2004).

22. K Doya, Bifurcations in the learning of recurrent neural networks in [Proceedings] 1992 IEEE International Symposium on Circuits and Systems. Vol. 6, pp. 2777–2780 vol.6 (1992).

23. AH Ribeiro, K Tiels, J Umenberger, T Schön, LA Aguirre, On the smoothness of nonlinear system identification. Automatica 121, 109158 (2020).

24. R Pascanu, T Mikolov, Y Bengio, On the difficulty of training recurrent neural networks in Proceedings of the 30th International Conference on Machine Learning, ICML’13. (Atlanta, GA, USA), Vol. 28, pp. 1310–1318 (2013).

25. J Keener, J Sneyd, Mathematical Physiology. (Springer, New York, NY) Vol. 8/1, 2 edition, (2009).

26. AL Hodgkin, AF Huxley, A quantitative description of membrane current and its application to conduction and excitation in nerve. The J. Physiol. 117, 500–544 (1952).

27. S Boyd, L Chua, Fading memory and the problem of approximating nonlinear operators with Volterra series. IEEE Transactions on Circuits Syst. 32, 1150–1161 (1985).

28. IW Sandberg, Uniform approximation and the circle criterion. IEEE Transactions on Autom. Control. 38, 1450–1458 (1993).

29. J Golowasch, E Marder, Ionic currents of the lateral pyloric neuron of the stomatogastric ganglion of the crab. J. Neurophysiol. 67, 318–331 (1992).

30. AL Taylor, JM Goaillard, E Marder, How Multiple Conductances Determine Electrophysiological Properties in a Multicompartment Model. J. Neurosci. 29, 5573–5586 (2009).

31. EM Izhikevich, Dynamical Systems in Neuroscience. (MIT Press, Cambridge, MA), (2007).

32. GB Ermentrout, DH Terman, Mathematical Foundations of Neuroscience. (Springer, New York), (2010).

33. G Drion, A Franci, J Dethier, R Sepulchre, Dynamic Input Conductances Shape Neuronal Spiking. eneuro 2, ENEURO.0031–14.2015 (2015).

34. WV Geit, ED Schutter, P Achard, Automated neuron model optimization techniques: A review. Biol. Cybern. 99, 241–251 (2008).

35. M Almog, A Korngreen, Is realistic neuronal modeling realistic? J. Neurophysiol. 116, 2180–2209 (2016).

36. DA McCormick, Y Shu, Y Yu, Hodgkin and Huxley model — still standing? Nature 445, E1–E2 (2007).

37. E Marder, D Bucher, Understanding Circuit Dynamics Using the Stomatogastric Nervous System of Lobsters and Crabs. Annu. Rev. Physiol. 69, 291–316 (2007).

38. DM Blitz, MP Nusbaum, State-Dependent Presynaptic Inhibition Regulates Central Pattern Generator Feedback to Descending Inputs. J. Neurosci. 28, 9564–9574 (2008).

39. E Marder, GJ Gutierrez, MP Nusbaum, Complicating connectomes: Electrical coupling creates parallel pathways and degenerate circuit mechanisms. Dev. Neurobiol. 77, 597–609 (2017).

40. B Johnson, J Peck, R Harris-Warrick, Differential modulation of chemical and electrical components of mixed synapses in the lobster stomatogastric ganglion. J. Comp. Physiol. A 175, 233–249 (1994).

41. M Innes, Flux: Elegant machine learning with julia. J. Open Source Softw. (2018).

42. Z Liu, J Golowasch, E Marder, LF Abbott, A Model Neuron with Activity-Dependent Conductances Regulated by Multiple Calcium Sensors. J. Neurosci. 18, 2309–2320 (1998).

43. KJ Aström, RM Murray, Feedback Systems: An Introduction for Scientists and Engineers. (Princeton University Press, Princeton), (2008).

44. PSC Heuberger, PMJ van den Hof, B Wahlberg, Modelling and Identification with Rational Orthogonal Basis Functions. (Springer, London), (2005).

45. D Nguyen, B Widrow, Improving the learning speed of 2-layer neural networks by choosing initial values of the adaptive weights in 1990 IJCNN International Joint Conference on Neural Networks. pp. 21–26 vol.3 (1990).

46. L Farina, S Rinaldi, Positive Linear Systems: Theory and Applications. (John Wiley & Sons), (2000).

47. HDI Abarbanel, DR Creveling, JM Jeanne, Estimation of parameters in nonlinear systems using balanced synchronization. Phys. Rev. E 77 (2008).

48. AH Ribeiro, K Tiels, LA Aguirre, T Schön, Beyond exploding and vanishing gradients: Analysing RNN training using attractors and smoothness in Proceedings of the Twenty Third International Conference on Artificial Intelligence and Statistics. (PMLR), pp. 2370–2380 (2020).

49. L Ljung, System Identification: Theory for the User. (Prentice Hall PTR, Upper Saddle River, NJ), (1999).

50. GJ Gutierrez, RG Grashow, Cancer Borealis Stomatogastric Nervous System Dissection. J. Vis. Exp. : JoVE p. 1207 (2009).

## References

1. D Nguyen, B Widrow, Improving the learning speed of 2-layer neural networks by choosing initial values of the adaptive weights in 1990 IJCNN International Joint Conference on Neural Networks. pp. 21–26 vol.3 (1990).

2. L Farina, S Rinaldi, Positive Linear Systems: Theory and Applications. (John Wiley & Sons), (2000).

3. D Angeli, ED Sontag, Monotone control systems. IEEE Transactions on Autom. Control. 48, 1684–1698 (2003).

4. PSC Heuberger, PMJ van den Hof, B Wahlberg, Modelling and Identification with Rational Orthogonal Basis Functions. (Springer, London), (2005).

5. AH Ribeiro, K Tiels, J Umenberger, T Schön, LA Aguirre, On the smoothness of nonlinear system identification. Automatica 121, 109158 (2020).

6. AH Ribeiro, K Tiels, LA Aguirre, T Schön, Beyond exploding and vanishing gradients: Analysing RNN training using attractors and smoothness in Proceedings of the Twenty Third International Conference on Artificial Intelligence and Statistics. (PMLR), pp. 2370–2380 (2020).

7. Z Liu, J Golowasch, E Marder, LF Abbott, A Model Neuron with Activity-Dependent Conductances Regulated by Multiple Calcium Sensors. J. Neurosci. 18, 2309–2320 (1998).

8. AA Prinz, CP Billimoria, E Marder, Alternative to Hand-Tuning Conductance-Based Models: Construction and Analysis of Databases of Model Neurons. J. Neurophysiol. 90, 3998–4015 (2003).

